# Realism and resolution: How AI-predicted protein structures differ from experimental structures

**DOI:** 10.64898/2026.11.06.764824

**Authors:** Rob van der Kant, Javier Delgado Blanco, Luis Serrano, Joost Schymkowitz, Frederic Rousseau

**Affiliations:** Switch Laboratory, VIB Center for Neuroscience Leuven, Herestraat 49, 3000 Leuven, Belgium; Switch Laboratory, Department of Cellular and Molecular Medicine, KU Leuven, Herestraat 49, 3000 Leuven, Belgium; Switch Laboratory, VIB Center for AI and Computational Biology, Leuven, Herestraat 49, 3000 Leuven, Belgium; Centre for Genomic Regulation (CRG), The Barcelona Institute for Science and Technology, Dr. Aiguader 88, 08003 Barcelona, Spain; Universitat Pompeu Fabra (UPF), Barcelona, Spain; ICREA, Pg. Lluis Companys 23, 08010 Barcelona, Spain

## Abstract

Protein structure predictors can recover protein folds with remarkable accuracy, and their models are increasingly used as if they were experimental structures. Yet even for a confidently predicted fold, it remains difficult to judge how faithfully the model reproduces protein geometry. We introduce WhatRes, which assesses a structure from its coordinates on two complementary axes. Apparent resolution estimates the experimental resolution its local geometry most closely resembles. Realism measures how closely that geometry matches experimental structures at the same apparent resolution. Predicted models can score well on apparent resolution while showing systematically less variation in bond lengths and angles than is observed in high-resolution X-ray structures. WhatRes reveals this over-idealisation even in high-confidence models.

## Introduction

Protein structure prediction has outrun structure validation. AlphaFold2^1^, AlphaFold3^2^ and the open Boltz models^3^ now generate atomic models that are used, in practice, as if they were experimental, for stability and mutation-effect prediction, docking and design. Yet the question a practitioner actually faces, “is this model good enough for the calculation I want to run?” has no established answer. The confidence scores that ship with the predictors (pLDDT, ranking scores) report expected coordinate accuracy^1^, not physical validity. Direct comparisons with experimental density have shown that even high-confidence AlphaFold predictions can contain substantial local and global errors^4^, but such comparisons require experimental data and so cannot provide a general validation strategy for the majority of predicted structures.

An established way to assess a structure is to examine its geometry. MolProbity^5,6^ checks for atomic clashes and unusual side-chain and backbone conformations, then combines these checks into a single score. This score is expressed on a scale resembling X-ray resolution, indicating the resolution at which an experimental structure would typically have similar geometric quality. ResProx (Resolution-by-Proxy)^7^ estimates this equivalent resolution directly from approximately 25 features of the atomic coordinates and can be applied to both experimental and predicted structures. Other methods, including VoroMQA^8^, ProQ^9^, QMEAN^10^ and DeepUMQA-X^11^, also use structural features to estimate model quality, usually reporting a score without physical units. Web servers bring several of these measures together to help users assess experimental and predicted structures^12^. These approaches establish that a model’s coordinates contain information about its quality, even when experimental data are unavailable.

The classical geometric metrics are, moreover, losing power exactly where AI and modern cryo-EM models need them. Bond lengths and angles are always restrained during refinement, and at low resolution refinement programs can also restrain Ramachandran, rotamer and clash terms, so such structures “score much better on validation than they really are”^13^, and at the 2.5-4 Å resolutions typical of cryo-EM the coordinate-only metrics decorrelate from true backbone quality^14^. The recommended remedy is a different kind of check. CaBLAM^13^ eg, assesses backbone conformation from virtual dihedral angles built using only the Cα and carbonyl-oxygen positions; because it depends on neither side chains nor the specific quantities that refinement restrains, it flags mis-oriented peptide planes and other backbone errors that the standard metrics miss, including at the low resolutions where carbonyl oxygens are themselves poorly resolved.

Interpreting a resolution-equivalent score as a complete measure of model quality, however, makes an important assumption. Experimental resolution describes the spatial detail supported by diffraction data or a reconstructed density map, while the precision of the atomic coordinates also depends on data quality and refinement^15,16^. Geometry-based methods exploit the empirical relationship between resolution and stereochemistry to express model quality on a resolution-like scale^6,7^. Yet this relationship reflects both the experimental information and the geometric restraints used during refinement^17,18^. Its usefulness therefore depends on whether a model follows the geometric patterns represented in that experimental reference. There is no guarantee that predicted structures follow the same patterns. The AlphaFold3 study, for example, reports residual ligand chirality errors and occasional severe atomic clashes despite its substantial gains in prediction accuracy^2^. More generally, a favourable resolution-equivalent score does not establish that a model reproduces the full geometric variation found in experimental structures. This motivates two complementary questions: what experimental resolution does a model’s geometry resemble, and how closely does it follow the geometric distributions observed at that resolution?

The distinction between coordinate accuracy and physical validity is already evident in protein-ligand prediction. PoseBusters^19^ showed that deep-learning docking methods can place ligands close to their experimental positions while producing incorrect stereochemistry, distorted bond geometry or steric clashes. Such limitations also occur in newer predictors: the AlphaFold3 study reported ligand chirality violations in 4.4% of predictions in its PoseBusters benchmark, despite explicitly penalising these errors during model ranking^2^. AlphaFold2 illustrates another aspect of this distinction: its AMBER relaxation removes stereochemical violations without improving coordinate accuracy as measured by GDT or lDDT-Cα^1^. Good stereochemistry and accurate coordinates are therefore complementary requirements. Beyond detecting overt violations, this raises a further question: does apparently well-formed geometry reproduce the variation observed in experimental structures?

When a predicted model looks confident, is its geometry actually that of a real structure? We address this question with WhatRes, a geometry-only framework that assesses structures along two complementary axes. Apparent resolution describes the experimental resolution a structure’s local geometry resembles; realism measures how closely that geometry follows the distributions observed in experimental structures of the same apparent resolution. Both measures use atomic coordinates alone and require no experimental metadata. AlphaFold3 and Boltz2 models on average resemble ∼2.5 Å X-ray structures in apparent resolution, yet score well below experimental structures in realism. Together, the two axes identify confidently predicted models whose geometry departs from experimental patterns in ways that a single resolution-equivalent score can conceal.

## Results

### A two-axis, geometry-only readout of structural quality

WhatRes operates on coordinates alone (Figure 1). For any structure, experimental or predicted, we compute 623 per-structure descriptors, implemented in our whatres-features program (Supplementary Table S1). These span covalent geometry (bond lengths and angles); backbone and side-chain dihedrals (φ/ψ/ω, χ1–χ4) and rotamer quality; chirality and peptide planarity (Cα chirality, Cβ deviation); restraint-robust backbone conformation (CaBLAM virtual dihedrals); all-atom clashes and packing; surface and cavity properties; and atom-type pairwise-distance distributions. B-factor and pLDDT are also read from the coordinate files but are excluded from both scores, since they report data quality and model confidence rather than geometry. Per-residue metrics are scored against reference distributions compiled from high-resolution X-ray structures (2.0 Å or better), converted to percentiles and “surprise” scores (the −log₂ of the two-tailed probability), and aggregated to the structure level.

**Figure 1.**
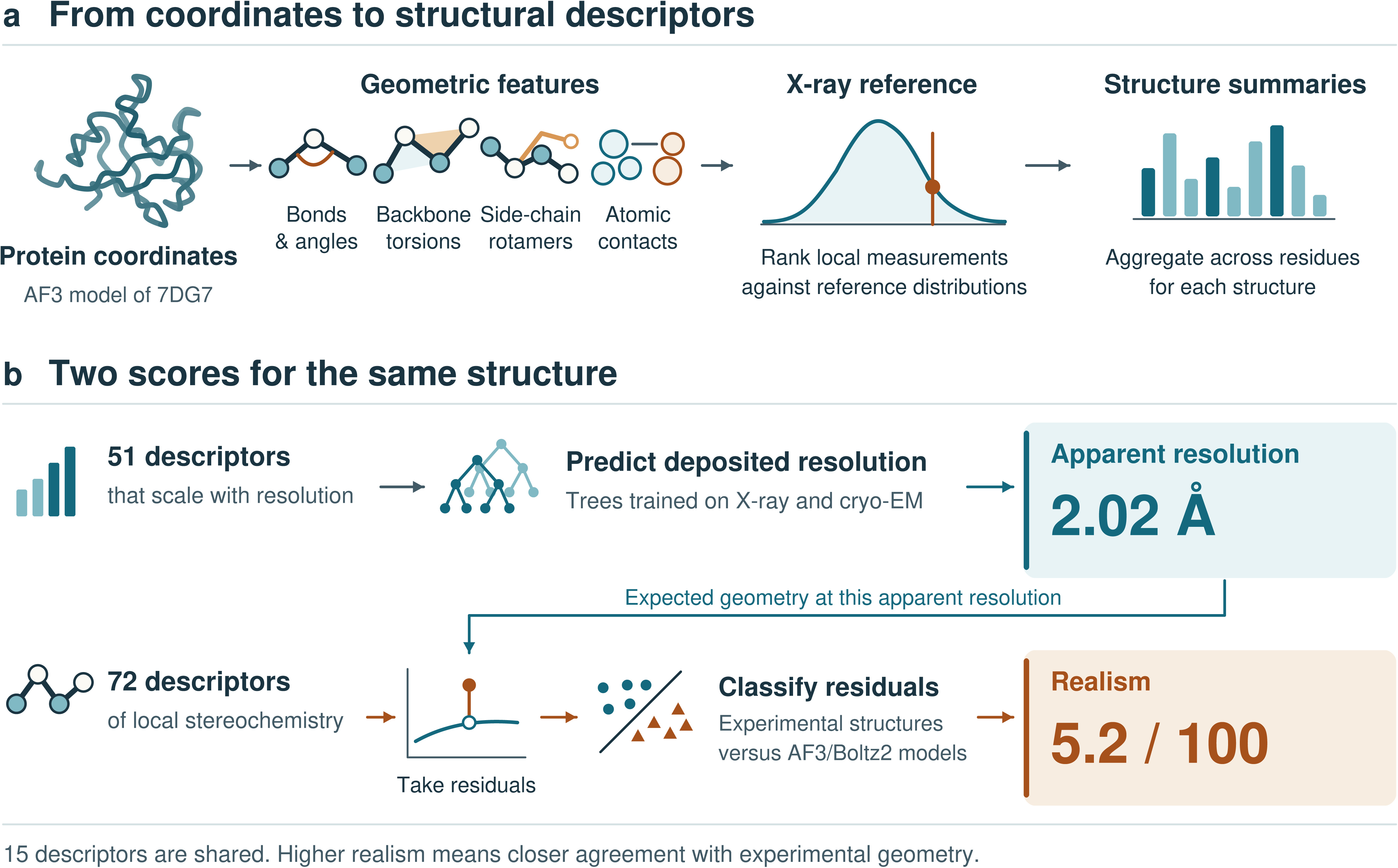
WhatRes estimates apparent resolution and geometric realism from protein coordinates. (a) The AlphaFold3 model of 7DG7 illustrates the workflow. WhatRes measures local geometry, including bond lengths and angles, backbone torsions, side-chain rotamers and atomic contacts. Local measurements are ranked against high-resolution X-ray reference distributions and summarised across residues to obtain structure-level descriptors. (b) A tree ensemble trained on X-ray and cryo-EM structures uses 58 descriptors to estimate apparent resolution. For each of 72 local stereochemical descriptors, WhatRes then subtracts the value expected for experimental structures at that apparent resolution. In the residual plot, the open circle marks the expected value and the filled circle marks the observed value. A classifier trained to distinguish experimental structures from AlphaFold3 and Boltz2 models uses these residuals to assign a realism score from 0 to 100. Higher scores indicate more experimental-like geometry. The two models share 19 descriptors. The 7DG7 prediction has an apparent resolution of 2.31 Å and a realism score of 6.7, illustrating how a model can resemble a well-resolved experimental structure in its resolution-sensitive features yet differ substantially in its local stereochemistry.

We first examine how each descriptor varies with deposited resolution across experimental X-ray and cryo-EM structures. Descriptors that vary systematically with resolution are combined by a feature-subsampled tree-ensemble regressor to estimate apparent resolution, expressed in Ångström. For realism, we use a panel of local geometric descriptors and remove the trend expected at each structure’s apparent resolution. A model trained to distinguish experimental from predicted structures then scores how closely the remaining geometry resembles that of experimental structures. The two scores use overlapping descriptor sets (19 are shared) and assess complementary aspects of geometry. Both are computed from coordinates alone and applied across our benchmark of experimental (X-ray, cryo-EM, NMR) and AI-predicted (AlphaFold2, AlphaFold3, Boltz2) structures.

### Predicted models occupy distinct regions of structure space

We first tested whether geometric descriptors alone distinguish structures produced by different methods. Using only the size-invariant geometric descriptors (with B-factor and pLDDT excluded, so the classifier is purely geometric), a random-forest classifier identifies the method of origin (X-ray, cryo-EM, NMR, AlphaFold2, AlphaFold3 or Boltz2) at 92% classification accuracy (Figure 2b). Predicted models are almost never mistaken for experimental structures (≤2% for each predictor), and the largest overlaps occur between X-ray and cryo-EM (5–10%, two density-based methods) and between AlphaFold3 and AlphaFold2 (10%, a shared lineage). This separating signal lies in low-variance directions of the descriptor space, so it is invisible in an unsupervised projection; a linear-discriminant analysis with 92% cross-validated accuracy reveals this separation (Figure 2a): experimental structures (X-ray and cryo-EM) occupy one region, the AlphaFold models a clearly distinct one and NMR a third, while Boltz2 falls between the AlphaFold and experimental clusters, closer to experimental structures in this projection than the AlphaFold models, consistent with its more experimental-like backbone geometry (Figure 2c-f). The classification and projection identify method-specific geometric patterns, consistent with reports of systematic differences in side-chain packing between predicted and experimental structures^20^.

**Figure 2.**
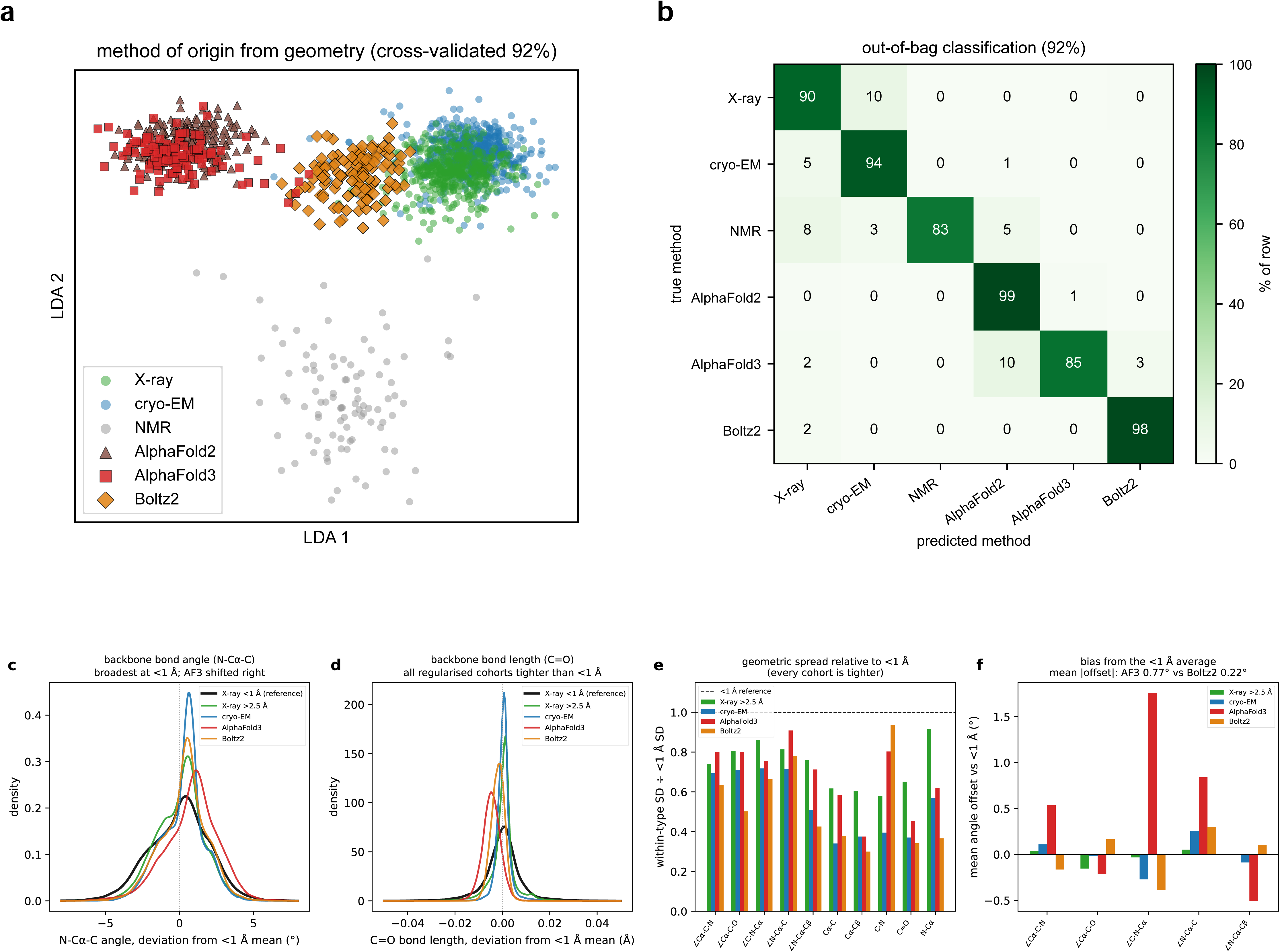
Geometry is method-specific and predicted models are over-regularised. (a) Supervised linear-discriminant projection of the size-invariant, geometry-only descriptor space (B-factor and pLDDT excluded), cross-validated accuracy 92%. Experimental structures (X-ray, green; cryo-EM, blue) occupy one region, AlphaFold models (AF2 brown, AF3 red) a distinct one, NMR (gray) a third, with Boltz2 (orange) intermediate. (b) Out-of-bag confusion matrix of a random-forest method-of-origin classifier (92% accuracy); values are percent of each true-method row. Over-regularisation of backbone geometry (c-f): raw backbone bond lengths and bond angles, computed directly from coordinates, for ultra-high-resolution X-ray structures (better than 1 Å; 49 structures, 9,890 residues), lower-resolution X-ray (worse than 2.5 Å), cryo-EM, AlphaFold3 and Boltz2, compared within residue type so that amino-acid composition and local conformation do not drive the comparison. (c) The backbone N-Cα-C bond angle and (d) the C=O bond length, each as the deviation from the ultra-high-resolution per-residue-type mean: ultra-high-resolution structures are broadest, every regularised cohort is tighter, and AlphaFold3 (c) is shifted to larger angles; the two modes of the N-Cα-C distribution are the α-helix and β-sheet sub-populations (the angle depends on φ/ψ; Supplementary Fig. 3). (e) Within-residue-type standard deviation of each backbone parameter as a fraction of the ultra-high-resolution value; every cohort lies below the dashed reference line at 1. (f) Mean offset of each backbone angle from the ultra-high-resolution average: AlphaFold3 is systematically displaced (mean absolute offset 0.77°, reaching +1.8° at C-N-Cα) whereas Boltz2 remains close to it (0.22°).

To understand this better, we compared raw backbone bond lengths and angles calculated directly from coordinates across five cohorts: ultra-high-resolution X-ray structures (better than 1 Å), where the diffraction data most directly determine local geometry; lower-resolution X-ray (worse than 2.5 Å); cryo-EM; and AlphaFold3 and Boltz2 (Figure 2c-f). Ultra-high-resolution structures show the broadest geometric distributions, and every other cohort is tighter in every parameter: within-residue-type standard deviations drop to 0.6 to 0.9 of the ultra-high-resolution value for lower-resolution X-ray structures and by up to roughly threefold for cryo-EM and the predictors (Figure 2c to e). This is the expected imprint of stereochemical restraints, which pull bond lengths and angles toward ideal targets when the experimental information is insufficient to define them^17,18,21,22^.

AlphaFold3 and Boltz2 also differ in their mean geometry: AlphaFold3’s backbone angles are systematically displaced from the ultra-high-resolution average (mean absolute offset 0.77° across five backbone angles, reaching +1.8° at C-N-Cα), whereas Boltz2 stays close to it (0.22°; Figure 2f). AlphaFold3 is therefore both over-regularised and biased, while Boltz2 is mainly over-regularised. This homogenisation extends to conformation-dependent geometry: the N-Cα-C backbone angle differs by about 2.9° between α-helix and β-sheet residues in ultra-high-resolution structures, reflecting its known dependence on backbone φ and ψ, and this helix-sheet separation is itself compressed to about 2.3° in every lower-resolution, cryo-EM and predicted cohort, with AlphaFold3’s angle bias largest in β-sheet (Supplementary Fig. 3).

### Apparent resolution: how much of the variation scales with resolution?

Several geometric differences in Figure 2 resemble the changes observed in experimental structures as data resolution worsens^17,23^. We therefore asked how well these resolution-dependent patterns could be used to estimate apparent resolution. We first ranked every descriptor by its Spearman correlation with deposited resolution across experimental structures and kept those that scale strongly with it (absolute Spearman above 0.3), excluding B-factor and pLDDT (which report data quality and model confidence, not geometry) and size-sensitive descriptors (which would let protein size, rather than geometry, drive the estimate); this left 58 descriptors. They come from several families, with atom-type pairwise distances and close-contact packing descriptors most numerous, together with backbone bond-length spread and rotamer-outlier level (Figure 3a), and each varies monotonically with resolution, though not all in the same direction: rotamer outliers become more frequent as resolution worsens, whereas the backbone carbonyl bond-length spread widens and close aliphatic-aliphatic contacts increase as resolution improves (Figure 3b).

**Figure 3.**
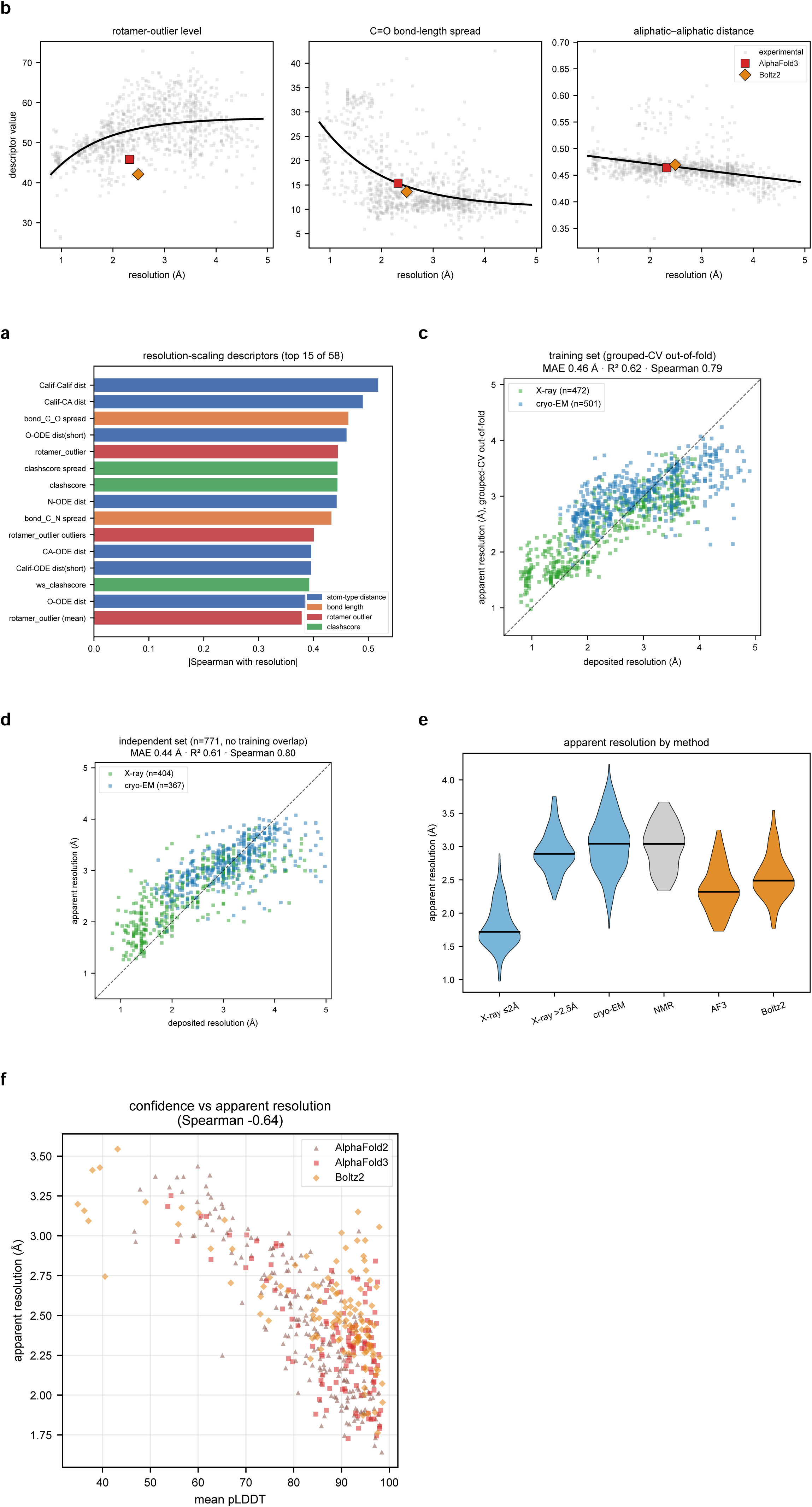
Apparent resolution. (a) The resolution-scaling descriptors, ranked by the absolute Spearman correlation of their deviation with deposited resolution and coloured by family; the 58 descriptors above the 0.3 threshold feed the apparent-resolution model. (b) Representative scaling descriptors against deposited resolution across experimental structures, with AlphaFold3 and Boltz2 overlaid at their apparent resolution, showing the models sit toward the high-resolution end. (c) Apparent resolution against deposited resolution for the experimental training set under grouped cross-validation (out-of-fold; 472 X-ray, 501 cryo-EM; MAE 0.46 Å, R² 0.62). (d) The same on an independent set of 771 deposited structures with no shared sequence cluster with the training set (MAE 0.44 Å, R² 0.61, Spearman 0.80). In c and d the dashed line is the identity; the below-unity best-fit slope reflects shrinkage of the estimator toward the training mean, not a calibration error, since the means coincide. (e) Apparent resolution by method of origin across the full benchmark (training structures by their out-of-fold predictions); the experimental classes order correctly from the highest-resolution X-ray structures to the lowest-resolution cryo-EM maps, and AlphaFold3 and Boltz2 reach favourable apparent resolutions (median about 2.3 and 2.5 Å). (f) Predictor confidence (pLDDT) against apparent resolution across all predicted models (Spearman −0.64). Descriptor-selection robustness (correlation-threshold sweep and random-subset control) is shown in Supplementary Fig. 4.

Using these descriptors, we trained an extremely randomised trees regressor to predict deposited resolution from experimental X-ray and cryo-EM structures. Feature subsampling reduced reliance on individual descriptors. We assessed performance using grouped cross-validation, holding MMseqs2 sequence clusters out together and repeating descriptor selection within each training fold. The model reproduced deposited resolution to a mean absolute error of 0.46 Å (R² = 0.62; 472 X-ray and 501 cryo-EM; Figure 3c). ResProx reports a lower error of 0.28 Å on a different, X-ray-only test set, so this comparison is indicative only. For a direct comparison, we evaluated the established MolProbity score on the same structures, which recovered the deposited resolution less accurately (error of 0.71 Å). Performance differed between experimental methods: X-ray tracks deposited resolution more tightly than cryo-EM (independent-set Spearman 0.82 versus 0.69; mean absolute error 0.45 and 0.42 Å), consistent with cryo-EM’s deposited resolution being a single global figure for a map whose local resolution varies. The estimate does not hinge on the exact descriptor set. Sweeping the selection threshold from 0.1 to 0.5, which changes the number of retained descriptors several-fold, leaves the cross-validated error stable through a threshold of 0.3 (0.46 to 0.47 Å), with a higher error at 0.4 (0.54 Å), while at 0.5 no descriptor passes in every training fold. The resolution-selected descriptors still outperform all twenty random draws of the same size (58 descriptors) from the size-invariant geometry panel (random error 0.54 to 0.66 Å; Supplementary Fig. 4), so the signal is spread across many resolution-scaling features rather than resting on a select few. Applied without retraining to an independent set of 771 deposited structures (404 X-ray and 367 cryo-EM, spanning 0.83 to 4.98 Å, with no shared MMseqs2 sequence cluster with the training set; three structures lacking cluster assignments were treated as singletons; see Methods), apparent resolution still tracks deposited resolution to a mean absolute error of 0.44 Å (R² = 0.61, Spearman 0.80; Figure 3d). Across the full benchmark, apparent resolution orders the experimental classes correctly, its median rising monotonically from ∼1.6 Å for the highest-resolution X-ray structures to ∼3.4 Å for the lowest-resolution cryo-EM maps (training structures scored by their out-of-fold predictions; Figure 3e).

Because apparent resolution is estimated from geometry alone, it can also be applied to predicted structures outside the experimental training domain. AlphaFold3 and Boltz2 receive favourable apparent resolutions, on average about 2.4 Å for AlphaFold3 and 2.5 Å for Boltz2, placing their resolution-dependent geometry within the range of good X-ray structures. The average, though, hides a wide spread: individual models range from about 1.7 Å, comparable to a high-resolution X-ray structure, to about 3.5 Å, more like a medium-resolution structure (5th to 95th percentile 1.9 to 3.1 Å). Predicted models are therefore far from uniform in these parameters, and apparent resolution distinguishes models whose geometry resembles experimental structures at different resolutions. Apparent resolution is also associated with predictor confidence: across all predicted models the two are substantially correlated (Spearman −0.64 between pLDDT and apparent resolution; Figure 3f), indicating that confidence and apparent resolution contain overlapping information. The association is incomplete, however, leaving variation in apparent resolution among confidently predicted models. We next examined geometric departures that remain after accounting for apparent resolution.

### Realism reveals variation among predicted models

Apparent resolution identifies the experimental resolution a structure’s geometry most closely resembles. To assess departures from experimental geometry beyond this resolution-dependent pattern, we developed a complementary realism score. We examined a panel of local stereochemical descriptors, including backbone and side-chain dihedrals, chirality, peptide planarity, cis-peptides, steric clashes, bond and angle geometry, and CaBLAM backbone measures. These descriptors differ in their dependence on experimental resolution, with some changing systematically with resolution and others remaining comparatively stable across experimental structures (Supplementary Fig. 4). Apparent resolution and realism are computed from overlapping geometric descriptors (19 of the 58 and 72 are shared). Realism assesses how closely local geometry resembles experimental structures after the resolution-dependent trend has been removed. The resolution-dependent descriptors, such as Cβ deviation and Cα chirality, scale with resolution and underlie apparent resolution (Figure 3). Predicted models also show less within-structure variation in backbone C=O bond length than high-resolution X-ray structures (median SD 0.004 Å for AlphaFold3 versus 0.007 Å for X-ray better than 2 Å; Figure 2d,e). Descriptors that vary little with experimental resolution reveal additional differences (Figure 4a). The most pronounced deviations occur in the longer side-chain dihedrals: χ3 and χ4 rotamer geometry varies little across experimental resolution, with comparable distributions in high- and low-resolution X-ray and cryo-EM structures, yet AlphaFold3 and Boltz2 depart systematically from this, showing higher dispersions. The shift extends across the distribution, including its median. Median χ3 surprise (−log₂ of its tail probability under the experimental reference; see Methods) is approximately 0.28 in experimental structures versus 0.68 in AlphaFold3. The differences remain after resolution correction (Cohen’s d 2.0 for χ3 and 1.7 for χ4). Thus, even typical predicted side chains fall in less probable regions of the experimental distribution. A further difference appears in the carbonyl bond length itself: the mean backbone C=O distance is essentially constant across the experimental resolution range examined (approximately 1.235 Å for X-ray at any resolution and for cryo-EM alike), although high-resolution X-ray structures show a broader distribution with a second population around 1.245 Å. AlphaFold3 and Boltz2 have shorter mean bond lengths, at 1.230 and 1.233 Å, close to the Engh and Huber restraint target of 1.231 Å ^21^, while the experimental cohorts lie consistently just above it (Figure 4a). The effect is selective: backbone ψ geometry is similarly insensitive to resolution but remains comparatively well reproduced by predicted models (Cohen’s d = −0.5; Figure 4a). These systematic differences expose a dimension of structural quality that apparent resolution alone cannot capture.

**Figure 4.**
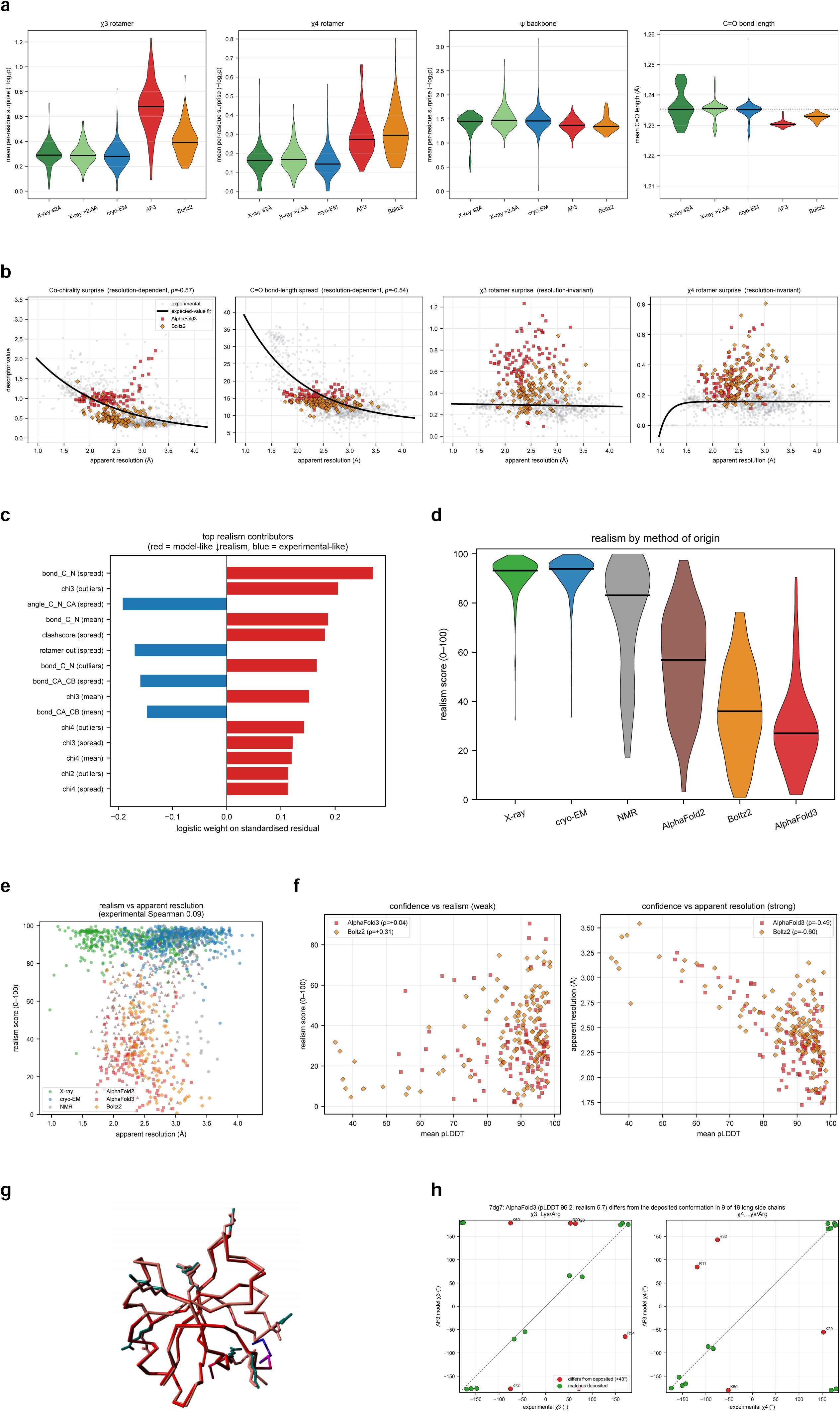
The realism axis. (a) Resolution-invariant descriptors that feed realism (χ3 and χ4 rotamer surprise, backbone ψ surprise, and the mean backbone C=O bond length), as per-structure distributions across X-ray (split at 2 Å), cryo-EM, AlphaFold3 and Boltz2: these are flat across experimental resolution, yet the predicted models are systematically offset in χ3 and χ4 and in the mean C=O bond length (centred short, on the Engh and Huber ideal of 1.231 Å) while reproducing ψ. (b) The residualisation step: representative descriptors against apparent resolution with the monotone experimental trend shown, AlphaFold3 and Boltz2 overlaid. (c) The strongest contributors to the realism classifier (signed logistic weights on the standardised residuals). (d) Realism score by method of origin (medians: X-ray 93, cryo-EM 94, NMR 83, AlphaFold2 57, Boltz2 36, AlphaFold3 27). (e) Realism against apparent resolution, showing the two axes are only weakly related (experimental Spearman 0.09). (f) Predictor confidence (pLDDT) against realism for AlphaFold3 and Boltz2 (weak; Spearman 0.04 and 0.31), in contrast to its strong association with apparent resolution. (g) A worked example of a confidently-predicted model that realism flags: the AlphaFold3 model of target 7dg7 (mean Cα pLDDT 96.2, realism 6.7) superposed on the experimental structure of the same target; the backbone is accurately predicted while nine of the nineteen lysine and arginine side chains differ from the deposited conformation. (h) The same model versus experiment, as the χ3 and χ4 angles of its Lys/Arg side chains; points off the diagonal (red, >40°) are side chains that differ from the deposited conformation (a valid but different rotamer), invisible to pLDDT and to a rotamer-outlier count (3% of residues) yet captured by realism.

To quantify this dimension, we developed a realism score that measures how closely a structure’s geometry resembles that of experimental structures after accounting for apparent resolution. The score uses 72 geometry-only descriptors (the local stereochemical panel above, summarised by each measure’s mean, dispersion and outlier fraction; Supplementary Table S3) and is calculated in two steps. First, using experimental X-ray and cryo-EM structures, we fit the expected value of each descriptor as a monotone (exponential-saturating) function of apparent resolution and subtract it from the observed value (Figure 4b). The resulting residual measures how much a structure deviates from experimental geometry at the same apparent resolution, removing the systematic contribution of resolution and placing structures of any apparent resolution on a common basis. Second, we train an L2-regularised logistic regression on the standardised residuals to distinguish experimental structures from predicted models (AlphaFold3 and Boltz2); the regularisation is deliberately strong (C = 0.003), so that the fitted probabilities retain a graded range rather than clustering at the extremes. The classifier identifies the combination of resolution-corrected deviations characteristic of predicted structures, with the shifts in the longer side-chain dihedrals among the strongest contributors (Figure 4c), and separates experimental from predicted structures almost perfectly in grouped, nested five-fold cross-validation (AUC 0.995). We report realism on a 0 to 100 scale as 100 × (1 − p), where p is the predicted probability of belonging to the model class, so experimental-like residual geometry scores near 100 and a model-like signature scores low, and we apply the fitted model unchanged to all held-out structures (NMR and AlphaFold2). The classifier is trained exclusively on the experimental-versus-predicted distinction, without using any downstream measure of structural quality. The resulting score is only weakly related to apparent resolution across experimental structures (Spearman ρ = 0.09; Figure 4e), with experimental structures receiving similarly high realism scores across the resolution range (Figure 4e and Supplementary Fig. 1). Yet it separates experimental and predicted structures sharply while revealing substantial variation among prediction methods (Figure 4d; Supplementary Fig. 2). X-ray and cryo-EM structures have median realism scores of 93 and 94, respectively, while NMR ensembles score somewhat lower (median 83). Predicted structures score substantially below these experimental references, with medians of 57 for AlphaFold2, 36 for Boltz2, and 27 for AlphaFold3, and broad interquartile ranges extending approximately from 18 to 72 (Figure 4d). Thus, although modern AI predictors generate models with favourable apparent resolution, their resolution-corrected geometry frequently differs substantially from that observed experimentally. Realism is only weakly associated with the predictors’ own confidence estimates (Spearman ρ = 0.04 for AlphaFold3 and 0.31 for Boltz2 between pLDDT and realism; Figure 4f left), in marked contrast to the stronger association between confidence and apparent resolution (Figure 4f right). High prediction confidence therefore does not imply experimental-like geometric realism. Together, apparent resolution and realism describe two complementary aspects of a structure’s geometry: what experimental resolution it resembles, and how closely it follows the geometric distributions observed at that resolution.

### Realism flags geometric departures missed by confidence and rotamer-outlier counts

The AlphaFold3 model of target 7dg7 illustrates how realism adds information beyond confidence and rotamer-outlier counts. It has a mean Cα pLDDT of 96.2, among the highest in the benchmark, and a realism score of 6.7 out of 100. Comparison with the experimental structure shows an accurately placed backbone, but 9 of its 19 lysine and arginine side chains differ from the deposited conformation, with χ3 or χ4 rotated by 90° to 160° (Figure 4g,h; Arg11, Arg23, Lys29, Arg32, Arg34, Arg54, Lys60, Lys72, Arg90). Eight of the nine are solvent-exposed (above 25% relative side-chain accessibility in the deposited structure, median 64%), placing most of these differences at conformationally less-constrained positions. Each differing side chain occupies a valid rotamer well, so these conformational differences need not appear as rotamer outliers. The overall rotamer-outlier fraction remains low, at 3% of residues, despite the differences from the deposited structure. Realism assigns a low score because the combined, resolution-corrected side-chain geometry departs from experimental patterns. It thus identifies a confidently predicted model with an unusual distribution of local geometry, even when conventional rotamer-outlier counts remain low.

### Model quality matters to energy functions, not to AI confidence scores

Because WhatRes scores any coordinate model, we asked whether model quality relates to the experimental success of de novo binders, using a published meta-analysis of 3,669 experimentally tested designs (394 binders) from 20 design campaigns against 15 targets, each re-predicted with AlphaFold3^24^. Design methods and success rates differ widely between campaigns, so all comparisons were made within campaigns, in the 13 campaigns with at least five binders and five non-binders. Neither realism nor apparent resolution separated binders from non-binders, whether computed on the design models or on the AlphaFold3 models (mean within-campaign AUROC 0.50 to 0.52), and adding them to the AlphaFold3 interface score ipSAE (mean AUROC 0.80) left its cross-validated performance unchanged (0.76; Supplementary Fig. 7a). Model quality did matter, however, for energy functions evaluated on the models. In the six campaigns where both good (realism ≥ 20 and apparent resolution ≤ 3.0 Å) and poor AlphaFold3 complexes included at least three binders and three non-binders, the FoldX interaction energy^25,26^ separated binders from non-binders better on good models in five, by a mean AUROC of 0.22 (p = 0.014; Supplementary Fig. 7b). The Rosetta interface energy gave a similar result (0.21, p = 0.012). The quality of a predicted model thus has no bearing on the confidence scores of the method that produced it, but it does affect an energy function computed on it.

## Discussion

WhatRes assesses protein geometry from coordinates alone along two complementary axes. Apparent resolution estimates deposited resolution with a mean absolute error of 0.46 Å in held-out experimental structures and 0.44 Å in an independent set spanning 0.83 to 4.98 Å. On the 973 structures scored by both methods, grouped cross-validation gave mean absolute errors of 0.46 Å for WhatRes and 0.70 Å for MolProbity. ResProx reports 0.28 Å on a different, X-ray-only test set. Realism measures departures from experimental geometry after accounting for apparent resolution, with which it is only weakly associated across experimental structures (rank correlation 0.09). Experimental structures score highly on realism (medians 93 for X-ray and 94 for cryo-EM), followed by NMR (83) and predicted models (57 for AlphaFold2, 36 for Boltz2 and 27 for AlphaFold3). Geometric descriptors also identify the method of origin with 92% accuracy. These results show that similar apparent resolutions can coexist with distinct geometric patterns.

Resolution-equivalent measures such as MolProbity and ResProx exploit the relationship between experimental resolution and model stereochemistry^17,23^. Our results show why this useful summary does not fully describe predicted structures. Models can have apparent resolutions around 2.5 Å, comparable to good X-ray structures, while departing systematically from the geometric distributions observed experimentally. Modern refinement and structure prediction can both produce tightly regularised geometry, making conventional measures of stereochemical quality harder to interpret. Realism addresses this limitation by assessing the geometry that remains after its resolution-dependent component has been removed.

The central finding is that predicted structures systematically under-represent the geometric variation observed experimentally. The comparison with ultra-high-resolution crystallography (better than 1 Å), where the data most directly determine local geometry, makes this particularly clear (Figure 2c-f). Every lower-resolution, cryo-EM and predicted cohort has narrower backbone bond-length and angle distributions than this reference. AlphaFold3 also shifts its mean backbone angles away from the ultra-high-resolution values, whereas Boltz2 shows much smaller shifts, consistent with its higher realism. The carbonyl bond provides a striking example: AlphaFold3 and Boltz2 have mean backbone C=O lengths near the Engh and Huber target of 1.231 Å, below the experimental mean of 1.235 Å. The predicted distributions therefore differ in both their spread and their centre.

The longer side-chain dihedrals show a further systematic departure. The χ3 and χ4 distributions shift toward less probable regions of the experimental reference, extending to typical predicted side chains, while backbone ψ remains comparatively well reproduced. Recent residue-level comparisons of AlphaFold-predicted and experimental side chains report the same deterioration toward the longer dihedrals, particularly χ3 and χ4 ^27^; our analysis detects this discrepancy from a predicted structure alone, without a matching experimental structure. Together, these findings suggest that predictors favour common geometric values while under-representing some of the variation present in experimental structures. Realism makes these distributional differences measurable after accounting for apparent resolution.

Predictor confidence provides limited information about these geometric departures. pLDDT is strongly associated with apparent resolution (Spearman −0.64) but only weakly with realism (0.04 for AlphaFold3 and 0.31 for Boltz2). An over-idealised model can therefore have both high confidence and favourable apparent resolution. Realism provides an independent geometric assessment. Although trained on the experimental-versus-AlphaFold3/Boltz2 contrast, it was also applied unchanged to NMR and AlphaFold2 structures, extending the assessment to structure classes not used in training.

Realism measures resemblance to experimental geometry, which provides a practical reference but does not establish physical correctness in every case. The carbonyl offsets are small in absolute terms (about 0.005 Å, comparable to coordinate uncertainty), yet their consistency across distributions and resolution ranges makes them detectable statistically. The per-residue reference distributions come from high-resolution X-ray structures, while the resolution-dependent trends are fitted to X-ray and cryo-EM structures. The classifier was trained against current predictors (AlphaFold3 and Boltz2), so its behaviour should be reassessed as prediction methods evolve. Its relevance to specific downstream modelling tasks also remains to be established. Score precision depends on protein size: the sampling uncertainty of realism scales approximately with the inverse square root of residue count (about 6 points for a 50-residue chain versus about 1 point for 400 residues or more) and is limited most by the sparse long side chains. Scores for small proteins therefore carry greater uncertainty (Supplementary Fig. 6).

This result has clear limitations. The benchmark compiles outcomes from many studies and assays, so some binding calls are probably wrong; such label noise weakens a signal rather than creating one. The comparison nonetheless rests on six campaigns, most with four or five binders among the good models. The largest campaign showed no difference, and the effect depends on where the quality threshold is drawn: it is largest at a realism threshold of 20 to 25 and small at 10 to 15 (Supplementary Fig. 7c). The design models could not be tested this way, because their realism mainly reflects the pipeline that produced them. A prospective test, with thresholds fixed in advance and binding measured in a single assay, is needed before model quality can be recommended as a filter in design workflows that rank candidates by computed energy.

WhatRes offers a practical assessment requiring coordinates alone. It places experimental and predicted structures on an apparent-resolution scale and flags departures from experimental geometric patterns before a model is used. For users, the two axes provide complementary information about local geometry; for developers, the differences in carbonyl geometry, backbone angles and side-chain rotamer distributions identify specific targets for improvement. Extending this assessment to complexes, nucleic acids, bound ligands and conformational ensembles is a natural next step.

## Methods

WhatRes judges a protein structure from its atomic coordinates alone, with no experimental data such as diffraction intensities or a density map, and reports two complementary scores. Both are built from the same starting point. For every residue we compute a large set of local geometric quantities (bond lengths and angles, backbone and side-chain torsions, chirality, atomic clashes, packing and inter-atomic distances), and compare each measured value to a reference distribution: the distribution of that same quantity among residues of the same amino-acid type and degree of burial, compiled from 2,000 high-resolution experimental X-ray structures (2.0 Å or better, 975,112 residues). This comparison turns each residue’s value into a surprise, defined as the −log₂ of the two-tailed tail probability of that value in the reference, so that a perfectly typical value scores 0 and the score grows the further into the tails the value lies; we equivalently record the value’s percentile within the reference (0 to 100). The two scores do not use these per-residue values directly. Instead, for each geometric quantity we summarise its per-residue surprises and percentiles across the whole structure into three numbers: the level, the mean surprise over all residues; the spread, the standard deviation of the per-residue percentiles; and the outlier fraction, the proportion of residues falling below the 5th or above the 95th percentile of the reference. Each structure therefore becomes a single fixed-length set of 623 such descriptors, independent of how large the protein is, and both scores are computed from that per-structure fingerprint. Apparent resolution places the structure on the familiar Ångström scale, and is built in three steps. First, among experimental structures, which carry a known deposited resolution, we keep the descriptors whose values change systematically with resolution. Second, we train a tree-ensemble regressor (extremely randomised trees) to predict deposited resolution from those descriptors, so that it learns the quantitative mapping from geometry to resolution using the experimental structures as worked examples. Third, we apply the trained regressor to any structure, experimental or predicted; it returns the resolution whose characteristic geometry the structure most resembles. A model whose geometry looks like a typical 2 Å X-ray structure is therefore reported as about 2 Å, even though it has no diffraction data of its own. Realism captures what resolution cannot, because a structure can look well resolved yet still be geometrically unrealistic in ways no real structure shows at any resolution. It is also built in three steps. First, we take the descriptors that reflect local geometry. Second, we remove the part that merely tracks resolution, by subtracting from each descriptor the value expected at the structure’s apparent resolution (this expected value is fit on the same experimental X-ray and cryo-EM structures), leaving only the geometry that is unusual for that resolution. Third, we train a regularised logistic regression on this residual geometry to separate genuine experimental structures from AI-predicted models. Realism is reported on a 0 to 100 scale, with experimental-like geometry near 100 and model-like geometry low. Apparent resolution is evaluated on structures held separate from training. Realism is trained only on the experimental-versus-predicted contrast and evaluated by grouped, nested cross-validation (see below). For every structure used to train either model, the scores we report come from cross-validation folds that did not contain it, because a fitted model reproduces its own training structures. The sections below give the datasets, descriptors, models and validation in full.

### Datasets

Reference distributions for per-residue scoring were compiled from 2,000 high-resolution X-ray structures (2.0 Å or better), comprising 975,112 residues, with each descriptor’s reference built separately for every amino-acid type and burial class. The training set for apparent resolution comprised experimental structures with a deposited resolution (X-ray and cryo-EM), with the exact value parsed from the PDB header (REMARK 2); 973 structures carried a usable deposited resolution. An independent validation set of 771 further deposited structures (404 X-ray and 367 cryo-EM, 0.83 to 4.98 Å), selected from 941 candidates with deposited resolution (58 lacked computed descriptors, and 112 were removed for sharing an MMseqs2 sequence cluster with the training set or for lacking X-ray/cryo-EM resolution), was used to test transfer. Three validation and eighteen training structures carry no cluster assignment and are treated as singletons; excluding the three leaves the transfer error unchanged at 0.43 Å. The benchmark used throughout also included 95 solution NMR structures and 223 AlphaFold2 models from the AlphaFold Database^28^. For the AI comparison we used a leakage-stratified prediction set of 120 protein targets, each predicted with AlphaFold3 and with Boltz2, stratified by temporal cutoff (AlphaFold3 training cutoff 2021-09-30; Boltz2 2023-06-01) crossed with sequence and structural homology to the pre-cutoff PDB. Full accessions and per-set counts are given in Supplementary Table S2. These sets play three distinct roles, kept separate throughout: the 2,000 high-resolution X-ray structures provide the reference distributions for the per-residue percentiles and surprises; the experimental structures carrying a deposited resolution (X-ray and cryo-EM) calibrate the resolution dependence of the structure-level descriptors, both for apparent resolution and for the residualisation used by realism; and the experimental and predicted structures together train the realism classifier to learn the resolution-independent geometric signature. All X-ray training structures were deposited between 2023 and 2026, so the reference and training geometry reflect current refinement practice. Because crystallographic refinement has changed over time and can regularise geometry at a given resolution^29^, we tested whether refinement era confounds the scores by comparing X-ray structures of fixed resolution (2.0 to 2.5 Å) deposited before 1995 with those deposited from 1995 onward (deposition years 1982 to 2025). Apparent resolution is independent of era (median 2.65 versus 2.52 Å, p = 0.56), whereas realism is modestly lower for the more recent structures (pre-1995 median 99.5, 1995-onward 96.5; p < 0.001), an effect confined to the restraint-sensitive local-geometry terms and roughly an order of magnitude smaller than the experimental-versus-predicted separation on the realism axis (Supplementary Fig. 5).

### Coordinate descriptors

For every structure we computed the 623 per-structure descriptors from coordinates alone, using our whatres-features program. The descriptors fall into the families enumerated in Supplementary Table S1: covalent geometry (backbone and Cβ bond lengths and bond angles); backbone dihedrals (φ, ψ, ω) and a two-dimensional Ramachandran density; side-chain dihedrals (χ1 to χ4) and a rotamericity proxy; chirality and planarity (Cα improper chirality, Cβ deviation from the ideal position, peptide-bond planarity, cis-peptide detection); restraint-robust backbone conformation (CaBLAM Cα and carbonyl-O virtual dihedrals); all-atom clashes and packing (a MolProbity-style all-atom clashscore, re-implemented natively); surface and cavity descriptors (solvent accessibility, hydrophobic patches, charge clusters, internal cavities, packing density and void index); atom-type pairwise-distance distributions; and per-residue B-factor or pLDDT statistics. Geometric, chirality, CaBLAM^13^, clashscore, surface and distance descriptors are computed by our own C++ code, as are the backbone dihedrals and the side-chain burial that assigns each residue its reference class: the occupancy of the side-chain atoms by neighbouring heavy atoms (Gaussian-weighted atomic volumes within 6 Å)^26^, scaled between the occupancy of a fully exposed and a fully buried atom and divided into five classes. No energy function is used. The atom-type pairwise-distance descriptors derive from histograms of residue-residue contacts resolved by atom type (60 bins from 0 to 9 Å), summarised per atom-type pair into the short-range (under 4 Å) and mid-range (4 to 6 Å) contact content and a per-contact-normalised form.

### Reference distributions and surprise scoring

Each per-residue metric was scored against reference distributions stratified by amino-acid type and burial, with a separate two-dimensional grid for the Ramachandran density. For a given residue, a metric value was converted to a percentile rank within its reference cell and to a surprise score, defined as S = −log₂[2·min(F, 1 − F)], where F is the value’s position in the reference cumulative distribution for its cell. A value at the median scores 0, one at the 95th percentile −log₂(0.1) ≈ 3.3 bits, and one at the 99th percentile −log₂(0.02) ≈ 5.6 bits. Surprise therefore measures statistical extremeness relative to high-resolution experimental geometry rather than energetic improbability, so that values typical of high-resolution structures score near zero and unusual values score high. Per-residue percentiles and surprise scores were aggregated to the structure level as the mean, the standard deviation, the fraction of residues with an extreme percentile (below 5 or above 95), and the mean surprise. The percentile-spread descriptor (the standard deviation of a structure’s per-residue percentiles) measures how widely residues are dispersed across the reference rather than the magnitude of their geometric deviations; because percentiles are approximately uniform under the reference, a structure matching experimental geometry has a characteristic non-zero spread, whereas an unusually small spread indicates residues clustered near the reference median, as in an over-idealised model. We therefore do not treat a smaller spread as better geometry: the sign and weight of every summary are learned by the models, not assumed.

### Resolution screen and the two feature classes

To separate the descriptors into the two classes used by the two scores, we computed, across experimental structures with a deposited resolution, the Spearman correlation between each descriptor and resolution. Descriptors whose deviation increases systematically with resolution define the resolution-dependent class; descriptors that are flat with respect to resolution, that is, physical constraints respected at every resolution, define the resolution-invariant class.

### Apparent resolution

Apparent resolution was predicted with an extremely randomised trees regressor (ExtraTreesRegressor, scikit-learn; 600 trees, each split drawn from a random 30% subset of the descriptors) trained to predict deposited resolution from the 58 resolution-scaling descriptors. The random feature subsampling is deliberate: a single boosted model concentrated most of its weight on one descriptor (the per-structure spread of the backbone C=O bond length), so that a small geometric perturbation could swing the estimate by about 0.3 Å through that one feature; feature-subsampled trees, which also randomise the split thresholds, spread the reliance across correlated descriptors and halve this sensitivity at essentially no cost in accuracy. To prevent the estimate from being driven by protein size or by modelled B-factors rather than intrinsic geometry, size-extensive features (absolute Spearman correlation with residue count above 0.35, including residue count itself) and all B-factor or pLDDT features were withheld. Performance was assessed by grouped cross-validation, with folds defined by MMseqs2 sequence clusters^30^ (30% identity and 80% coverage) so that homologues are held out together, and with the descriptor selection repeated within each training fold, giving a mean absolute error of 0.46 Å (R² 0.62); applied without retraining to the independent 771-structure set, itself family-separated from the training set by the same clustering, it reached 0.44 Å (R² 0.61, Spearman 0.80). Because the fitted ensemble (fully grown trees without bootstrap resampling) reproduces the deposited resolution of its own training structures exactly, every apparent resolution we report for a training structure, in any figure or analysis, is its out-of-fold prediction. Because the predictor minimises squared error it behaves as a shrinkage estimator, compressing its output toward the training mean, so the best-fit slope of apparent against deposited resolution is below one (about 0.5 in training and on the independent set) while the two means coincide; the identity line is therefore the reference in Figure 3c,d, and accuracy is summarised by mean absolute error, R² and rank correlation rather than by the regression slope. As a baseline we computed the genuine MolProbity score on the same structures with the official MolProbity programs and reference data: the all-atom clashscore from Word’s probe on reduce-added hydrogens, together with rotamer-outlier and Ramachandran-favoured percentages from the Top8000 contour grids^31^, combined by the Chen et al. (2010) formula; this pipeline reproduces phenix.molprobity^32^ to within 0.1 MolProbity-score units on a reference structure. The MolProbity score recovered deposited resolution with a mean absolute error of 0.71 Å after a linear calibration to this cohort, fitted within the same grouped cross-validation folds, comparable to apparent resolution. The dedicated ResProx predictor reports 0.28 Å on its own, X-ray-only test set (a different set, so the comparison is indicative only) (its server was not reachable during this work, so we cite the published value rather than run a head-to-head). Apparent resolution was trained on both X-ray and cryo-EM structures rather than X-ray alone: cryo-EM deposited resolution is a single global figure for a map whose local resolution varies, which adds label noise (cross-validated R² 0.41 versus 0.71 for X-ray, Spearman 0.68 versus 0.87, and a shallower fitted slope, 0.34 versus 0.57), but including it calibrates apparent resolution across the 2.5 to 4 Å regime where most cryo-EM and predicted models fall, keeping them within the training domain rather than as pure extrapolation (mean absolute error on held-out cryo-EM 0.42 to 0.52 Å). Restricted to X-ray, where ResProx was benchmarked, our mean absolute error is 0.41 Å.

### Realism score

The realism score measures a structure’s similarity to the resolution-corrected geometric characteristics of experimental structures; operationally it flags geometry that departs from that of real experimental structures of the same apparent resolution, in particular geometry more idealised than any real structure. It thus quantifies experimental likeness rather than physical plausibility in the abstract. It is built from 72 geometry-only descriptors (Supplementary Table S3) spanning rotamers (χ1 to χ4) and a rotamericity proxy, backbone dihedrals (φ, ψ, ω), chirality and planarity, bond and angle statistics, the all-atom clashscore and CaBLAM backbone measures, each summarised by its mean surprise, dispersion and outlier fraction. To make the score independent of resolution, we first removed from each descriptor the part explained by apparent resolution: for every descriptor we fit its expected value as a monotone (exponential-saturating) function of apparent resolution across experimental structures and replaced the descriptor by its residual from that curve, so that a real structure of any resolution sits near zero by construction. We then trained an L2-regularised logistic regression (standardised features) to separate experimental structures from predicted models (AlphaFold3 and Boltz2) on these residuals; the penalty was set strong (C = 0.003) so that the fitted probability is graded rather than saturating. In grouped five-fold cross-validation (folds defined by MMseqs2 sequence clusters, each predicted model taking the cluster of its target, so that a target’s AlphaFold3 model, its Boltz2 model and its experimental structure share a fold), with the residualisation curves and the classifier refitted within every training fold, the classifier separates experimental from predicted structures almost perfectly (AUC 0.995). The residualisation uses the out-of-fold apparent resolution of each experimental training structure, the value any new structure would receive, and every realism score we report for a training structure is its out-of-fold score. Realism is reported on a 0 to 100 scale as 100 times one minus the model probability, so that experimental structures score near 100 and strongly over-idealised models score low, and equivalently as a 1 to 6 grade. Because each descriptor is residualised against apparent resolution before training, realism is only weakly correlated with apparent resolution in practice (Spearman 0.09 across experimental structures). The fitted model was applied unchanged to all held-out structures (NMR and AlphaFold2). Both scores use geometry-derived descriptors only.

### Orthogonality and method-of-origin

To test that resolution is a direction in descriptor space, we fitted a ridge regression of deposited resolution on the standardised descriptors and projected structures onto the unit resolution axis; the projection, computed out-of-fold under the same grouped cross-validation, correlated with deposited resolution at Spearman 0.72. Method of origin was classified from the size-invariant, geometry-only descriptors (B-factor and pLDDT excluded) with a random-forest classifier (scikit-learn, 500 trees), evaluated out-of-bag, distinguishing the six classes (X-ray, cryo-EM, NMR, AlphaFold2, AlphaFold3 and Boltz2) at 92% out-of-bag accuracy. The separating signal lies in low-variance directions of the descriptor space; to visualise it we used a cross-validated linear-discriminant projection (LinearDiscriminantAnalysis, eigen solver with automatic shrinkage; 92% cross-validated accuracy). Descriptors were quantile-transformed to an approximately normal distribution before both analyses. Each tree is grown on a bootstrap resample, so the out-of-bag estimate needs no separate held-out split; correlated descriptors were not collapsed into clusters, because each split draws on a random subset of features and the out-of-bag accuracy is insensitive to this redundancy, and the cross-validated discriminant projection confirms the separation is not carried by a single feature.

### De novo binder benchmark

We used the meta-analysis dataset of Overath et al.^24^ (Zenodo, 10.5281/zenodo.15722219; CC BY 4.0): 3,676 designs, of which 3,669 carry an experimental outcome, with the designers’ models, AlphaFold3 re-predictions of each complex, Rosetta interface energies and ipSAE. WhatRes scored each design model and the top-ranked AlphaFold3 model, as complex and as binder chain. FoldX AnalyseComplex^25,26^ computed binder-target interaction energies on the unrepaired complexes; the 137 complexes it could not analyse all belong to two pMHC campaigns outside every testable campaign. Campaigns (one source and one target) with at least five binders and five non-binders were analysed. Predictors were compared by the AUROC between binders and non-binders within each campaign, averaged over campaigns, with p values from permuting labels within campaigns. Added value over ipSAE was assessed by logistic regression with five-fold cross-validation within each campaign. For the energy comparison, AlphaFold3 complexes were classed as good when realism was at least 20 and apparent resolution at most 3.0 Å, thresholds set before the energy analyses. Campaigns with at least three binders and three non-binders among both good and poor models were compared, with p values from permuting good and poor labels within campaigns.

### Software

Descriptor computation used whatres-features (C++). Analyses used Python with scikit-learn^33^ (ExtraTreesRegressor for apparent resolution, a regularised LogisticRegression for realism, RandomForestClassifier and LinearDiscriminantAnalysis for method of origin, and Ridge for the resolution axis), scipy^34^, pandas^35^, numpy^36^ and matplotlib^37^. The MolProbity baseline used the official MolProbity programs: probe and reduce^38^ with the Top8000 rotamer and Ramachandran reference data^39^, through a cctbx/mmtbx installation^40^. Structure metadata were retrieved from the RCSB PDB^41^.

## Supporting information

WhatRes_Supplementary_Figures

WhatRes Suppl Table 2 datasets

WhatRes Suppl Table 1 Parameters

WhatRes Suppl Table 3 Realism parameters

## Data availability

The experimental structures analysed in this study are available from the Worldwide Protein Data Bank (wwPDB) under the accession codes listed in Supplementary Table S2. AlphaFold2 models were obtained from the AlphaFold Protein Structure Database. The AlphaFold3 and Boltz2 models generated for this study, the high-resolution reference distributions, and the per-structure apparent-resolution and realism scores are available at Zenodo (https://doi.org/10.5281/zenodo.24014748). Source data for the figures are provided with this paper. The de novo binder data analysed in Supplementary Fig. 7 are from Overath et al.^24^ (https://doi.org/10.5281/zenodo.15722219).

## Code availability

WhatRes is available at https://github.com/Schymkowitz/WhatRes under the BSD 3-Clause licence. It consists of whatres-features, a command-line program written in C++ (C++14; macOS and Linux) that computes the structural descriptors, and a Python package that assembles them and applies the two trained models (apparent resolution and realism), which are provided with it together with the high-resolution reference database. The atom parameters used for side-chain burial (per-atom volume and minimal and maximal occupancy^26^) are distributed with whatres-features as a precompiled library under a separate licence. Both scores are computed from coordinates alone and require no energy function. The code used to train the models and to produce the analyses, figures and tables of this study is available at https://github.com/Schymkowitz/WhatRes-paper. Both repositories are archived at Zenodo (https://doi.org/10.5281/zenodo.24015562; https://doi.org/10.5281/zenodo.24394888).

## Acknowledgements

The Switch Laboratory acknowledges support from the Flanders Institute for Biotechnology (VIB, grant C0401 to F.R. and J.S.) and KU Leuven. J.D.B. and L.S. acknowledge institutional support from the Generalitat de Catalunya through the CERCA programme and from the EMBL–CRG partnership. We thank Oscar Garrett (University of California, Davis) for suggesting the analysis of de novo binder outcomes.

## Author contributions

J.S. conceived the approach and wrote the software, with conceptual input from R.v.d.K., J.D.B., L.S. and F.R.; J.S. and R.v.d.K. designed the analyses and benchmarks and performed the analyses; R.v.d.K. and J.S. prepared the figures. J.D.B. contributed the predicted-model sets and descriptor tooling. L.S., F.R. and J.S. supervised the work and acquired funding. R.v.d.K., J.S. and F.R. wrote the manuscript with input from all authors. All authors read and approved the final manuscript.

## Use of artificial intelligence

The authors used large language model tools interactively to develop the analysis code and to draft and revise the manuscript. The authors directed this work and take full responsibility for the code, the analyses, the interpretation of the results, and the final text.

---

**Supplementary Figure 1.**
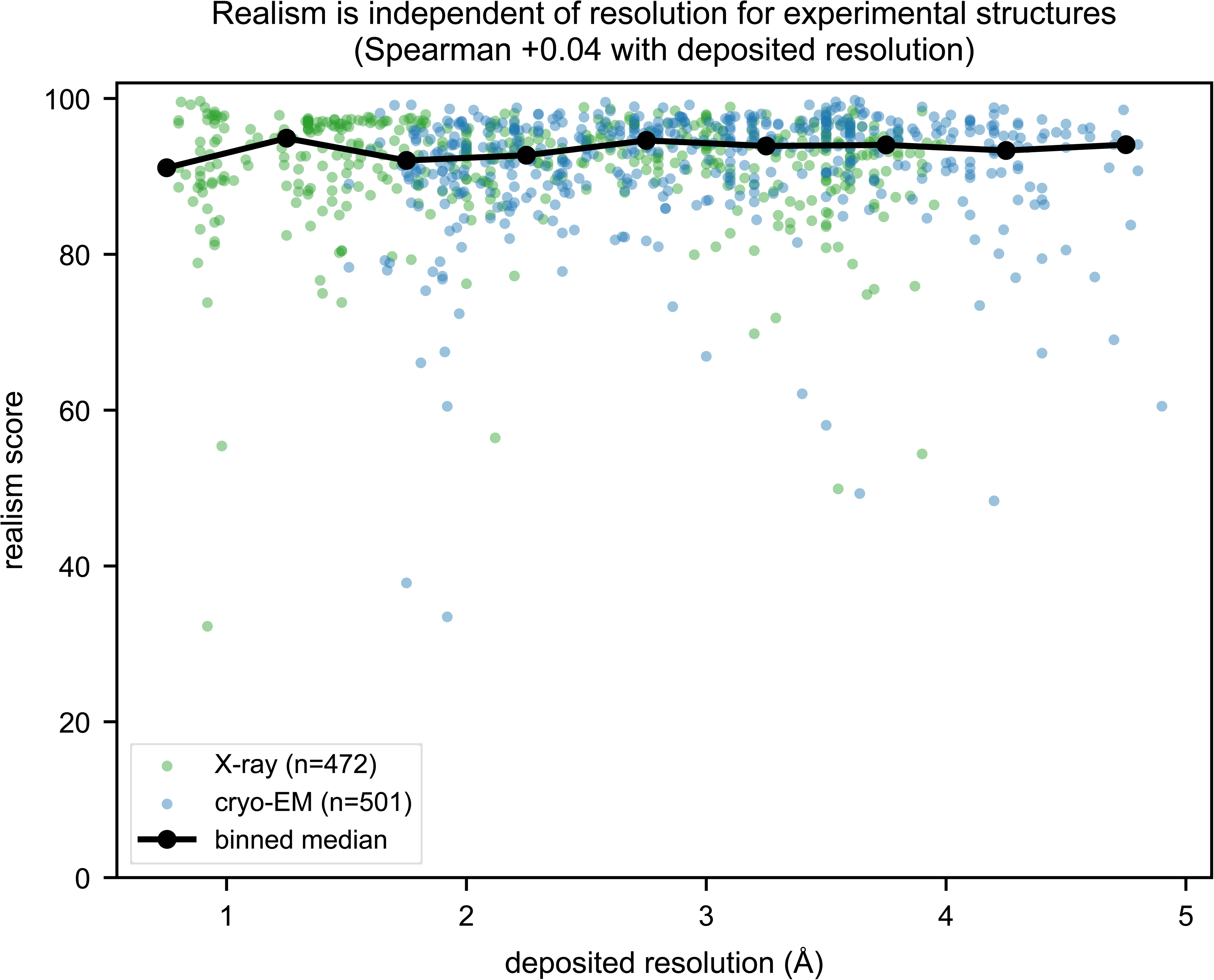
Realism is independent of resolution for experimental structures. Realism score against deposited resolution for experimental X-ray and cryo-EM structures, with the binned median overlaid. Realism remains high and approximately constant across the full resolution range (Spearman 0.04 with deposited resolution), confirming that the score reflects experimental-like geometry rather than the amount of experimental detail.

**Supplementary Figure 2.**
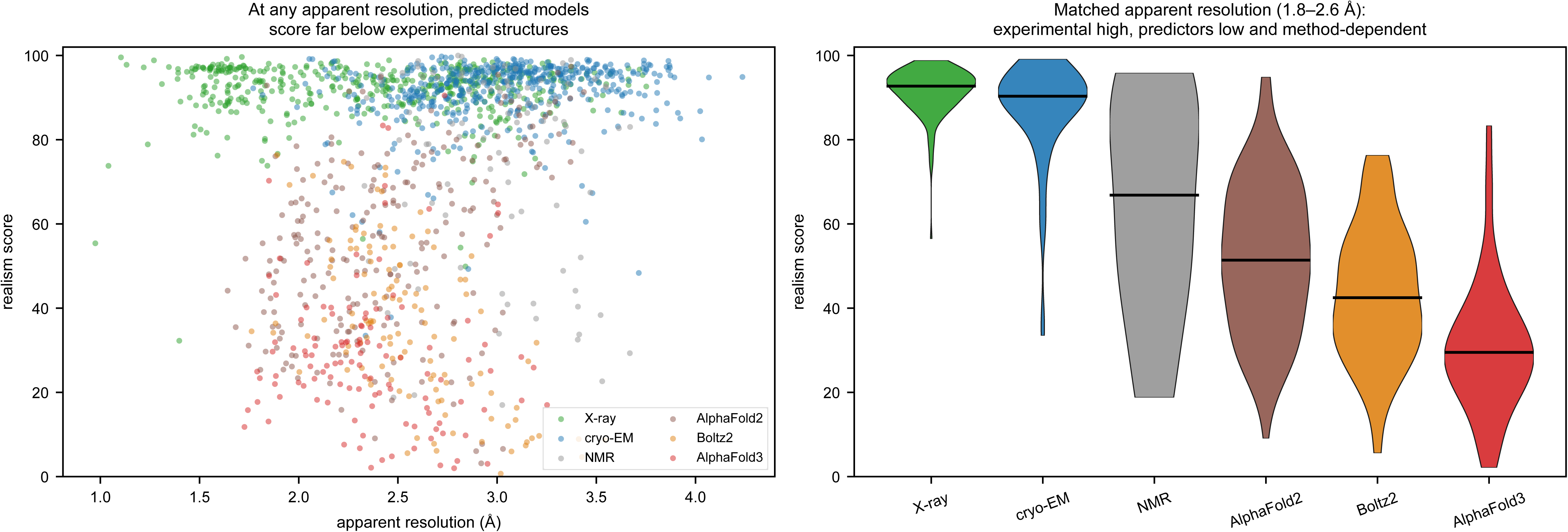
Predicted models score far below experimental structures at matched apparent resolution. (a) Realism against apparent resolution across the benchmark, coloured by method: experimental structures (X-ray, cryo-EM, NMR) remain high across the apparent-resolution range, whereas AlphaFold2, AlphaFold3 and Boltz2 sit well below them even at favourable apparent resolutions. (b) Realism distributions within a matched apparent-resolution band (1.8 to 2.6 Å): experimental structures score around 92 while the predictors score substantially lower and method-dependently (AlphaFold2 51, Boltz2 42, AlphaFold3 29), showing the separation is not a resolution artefact.

**Supplementary Figure 3.**
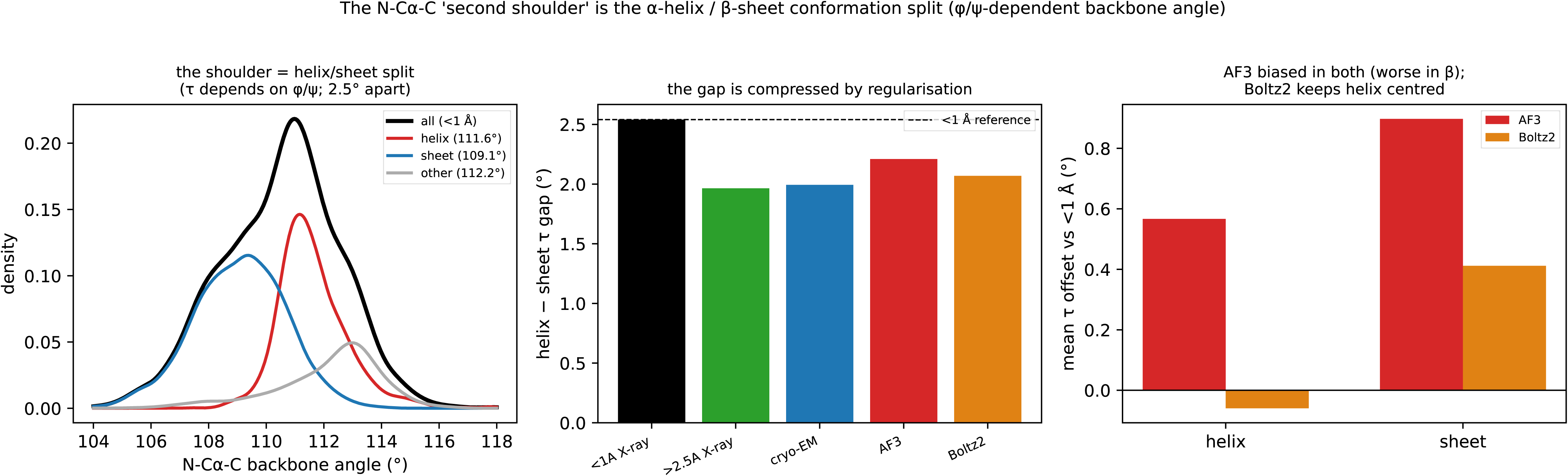
The N-Cα-C backbone angle by backbone conformation. The two modes of the N-Cα-C angle distribution in Figure 2c are α-helical and β residues. Conformation is assigned from φ and ψ (α: −100° < φ < −30° and −80° < ψ < −10°; β: φ < −90° and ψ > 90°), and Gly and Pro are excluded as in Figure 2c. (a) In ultra-high-resolution (<1 Å) X-ray structures the mean angle is 111.6° for α-helical and 108.8° for β residues, a difference of 2.9°, reflecting the dependence of the angle on φ and ψ. (b) This helix-sheet difference is compressed to 2.2 to 2.4° in the lower-resolution X-ray, cryo-EM and predicted cohorts (error bars, 95% confidence interval). (c) Offset of the mean angle from the ultra-high-resolution reference: AlphaFold3 is shifted to larger angles in both conformations and most in β residues (+0.6° helix, +1.2° sheet), whereas Boltz2 stays close to the reference in helices (−0.1°).

**Supplementary Figure 4.**
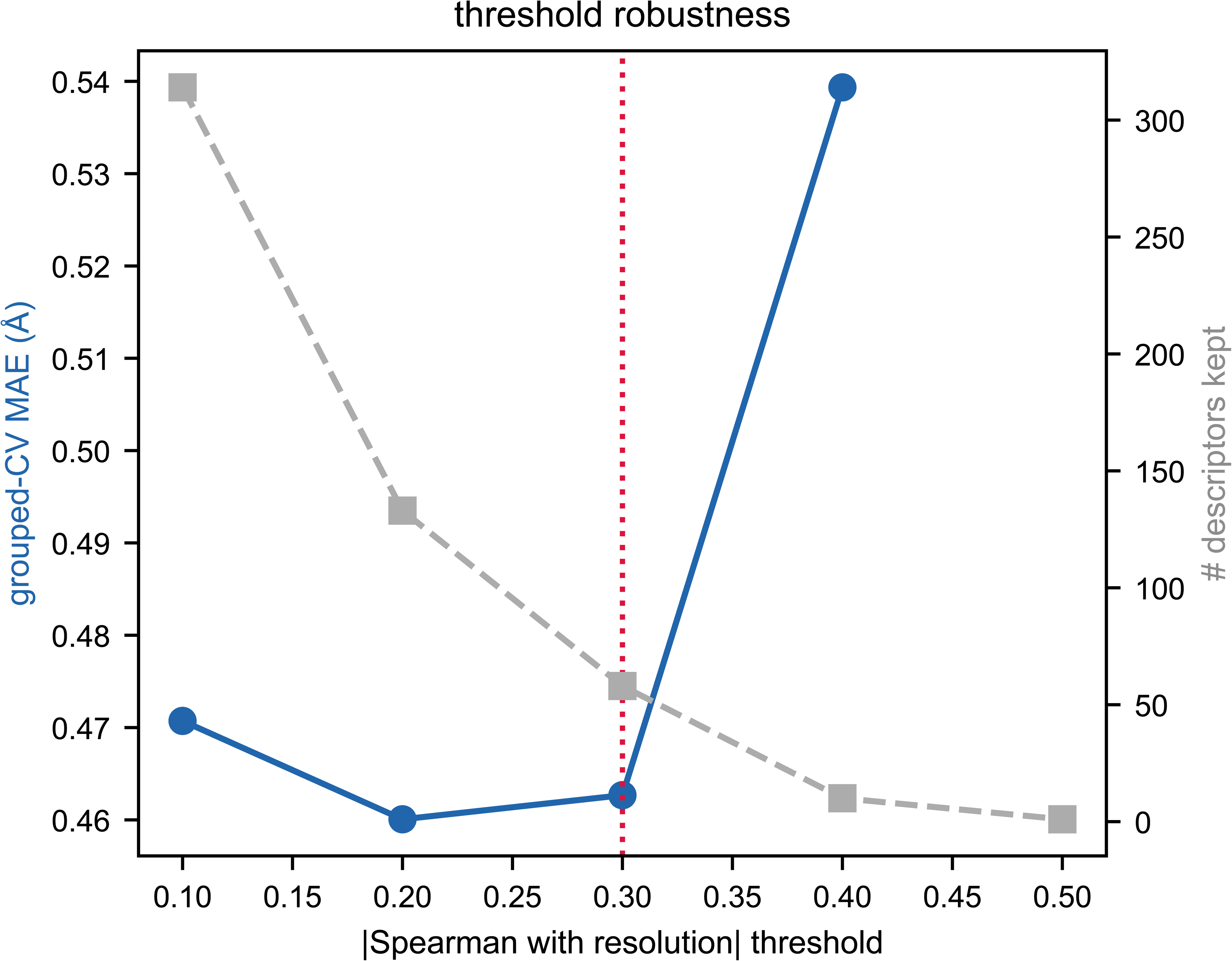
Robustness of the apparent-resolution descriptor selection. (a) Grouped cross-validated mean absolute error as the correlation threshold for retaining resolution-scaling descriptors is swept from 0.1 to 0.5; with the selection repeated within each training fold, the error is stable through 0.3 (0.46 to 0.47 Å) and rises from 0.4 (0.54 Å); at 0.5 no descriptor passes in every training fold. (b) The resolution-selected descriptors (selected within each training fold) versus twenty random 58-descriptor draws from the size-invariant geometry panel: the selected set outperforms every random draw (random error 0.54 to 0.66 Å), so the signal is spread across many resolution-scaling features.

**Supplementary Figure 5.**
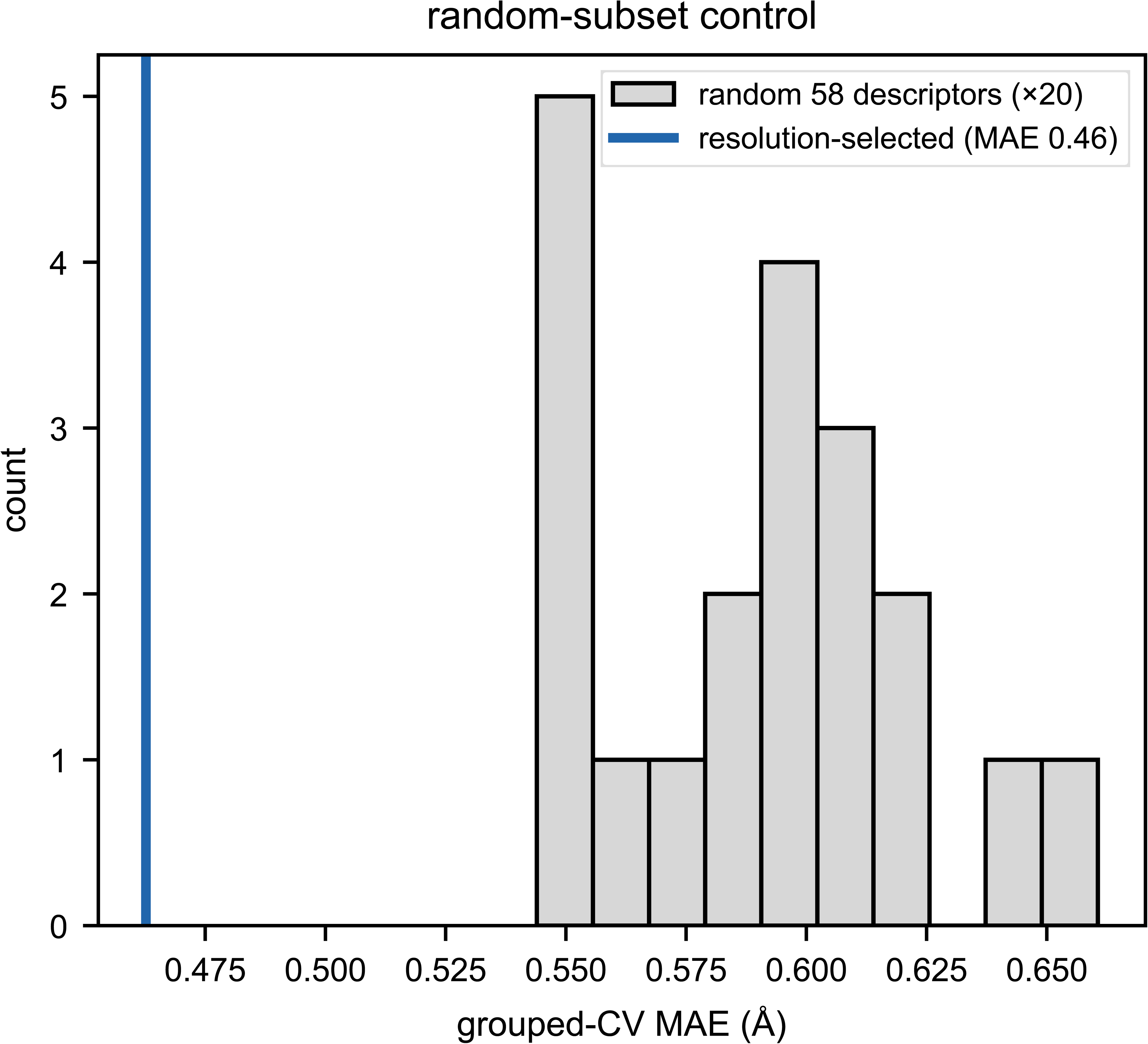
Apparent resolution and realism before and after 1995, at fixed resolution. Crystallographic refinement has been progressively automated and re-standardised since the late 1980s, with the adoption of maximum-likelihood targets^42^, improved stereochemical restraint libraries^21^ and, later, conformation-dependent geometry restraints^18^, which regularise model geometry at a given resolution, sometimes at the expense of experimentally supported variability^17^. We scored 100 X-ray structures at a fixed deposited resolution (2.0 to 2.5 Å): 35 deposited before 1995 and 65 from 1995 onward (deposition years 1982 to 2025), all re-featurised in a single pipeline run. (a) Apparent resolution is independent of era (median 2.65 versus 2.52 Å; Mann-Whitney p = 0.56; equal spread, Levene p = 0.14). (b) Realism is modestly lower for structures deposited from 1995 onward (median 96.5 for 1995-onward versus 99.5 for pre-1995, p < 0.001; Spearman with deposition year −0.40), with no significant difference in spread (standard deviation 12.6 versus 10.2, Levene p = 0.09). (c) The era-effect is confined to the restraint-sensitive local-geometry descriptors (backbone bond angles, peptide bond lengths, Cβ deviation, Cα chirality): structures deposited before 1995 carry more geometric scatter in exactly the quantities modern refinement homogenises. Apparent resolution is therefore unaffected by refinement era, while realism detects the known homogenisation of restrained local geometry; because this shifts modern structures slightly toward the idealised end, the effect is roughly an order of magnitude smaller than the experimental-versus-predicted separation on the realism axis (predicted medians 27 to 57 versus experimental 93 to 94) and leaves our conclusions unchanged, if anything making that contrast conservative.

**Supplementary Figure 6.**
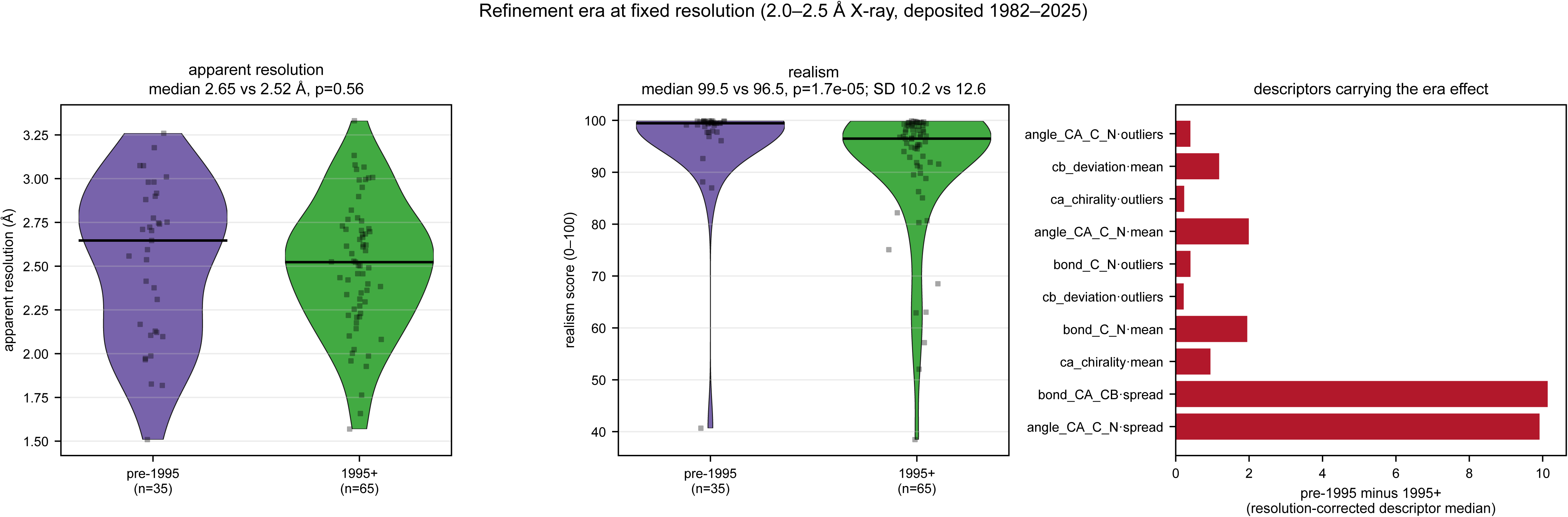
Protein size and score precision. Both scores are summarised per residue across the whole structure, making them size-independent in expectation but not in precision. (a) Realism against protein length across the benchmark, coloured by method: the median shows no strong length trend, but the spread widens for small proteins. (b) Sampling uncertainty measured within single large structures: for ten X-ray structures of 614 to 663 residues, realism was recomputed from random residue subsets of increasing size (100 resamples each); the score standard deviation falls approximately as the inverse square root of the subset size, from about 6 points at 50 residues to about 1 at 400 or more, with negligible bias. (c) The limiting sample size is set by the long side chains: χ3 is defined for about 23% of residues (Arg, Lys, Gln, Glu, Met) and χ4 for only about 11% (Arg, Lys), so a 100-residue protein provides roughly ten χ4 observations.

**Supplementary Figure 7.**
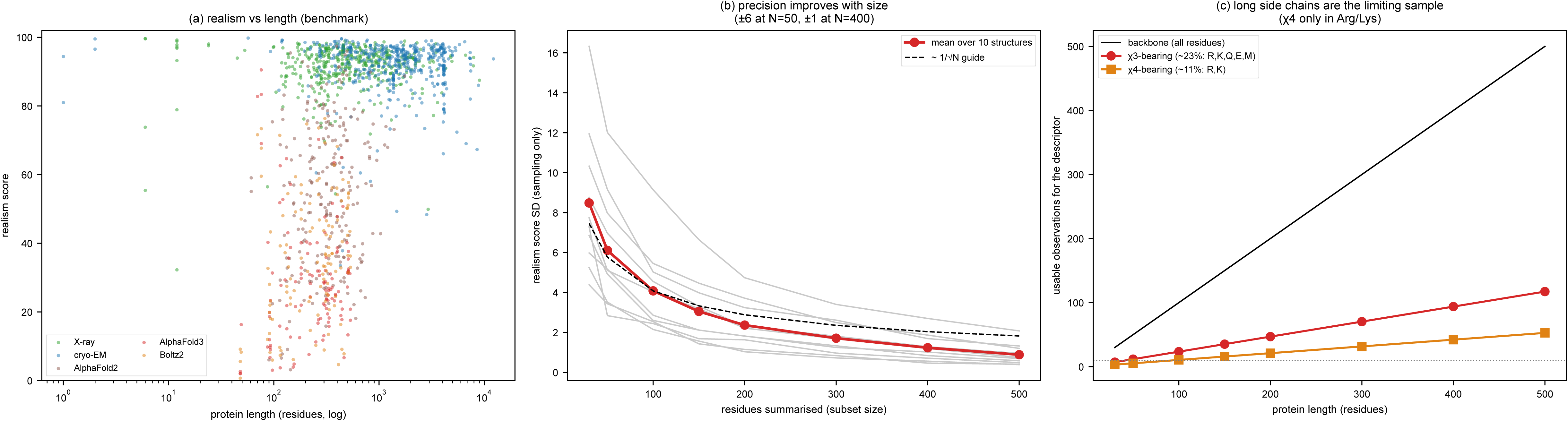
Model quality and the experimental success of de novo binders. Data from a meta-analysis of 3,669 experimentally tested designs (394 binders)^24^. (a) Within-campaign AUROC for separating binders from non-binders in the 13 campaigns with at least five binders and five non-binders (points, one per campaign; bars, means), for the AlphaFold3 interface score ipSAE, the Rosetta and FoldX interaction energies computed on the AlphaFold3 and design models, and WhatRes realism and apparent resolution of the same complexes. (b) AUROC of the interaction energy on AlphaFold3 complexes rated poor or good by WhatRes (good: realism ≥ 20 and apparent resolution ≤ 3.0 Å), for the 6 campaigns with at least three binders and three non-binders in both groups; lines join the same campaign. (c) Mean AUROC difference between good and poor models for the FoldX energy, against the realism threshold; n, number of campaigns that can be tested.

