## Supplementary material for "Realism and resolution: How AI-predicted protein structures differ from experimental structures": WhatRes_Supplementary_Figures

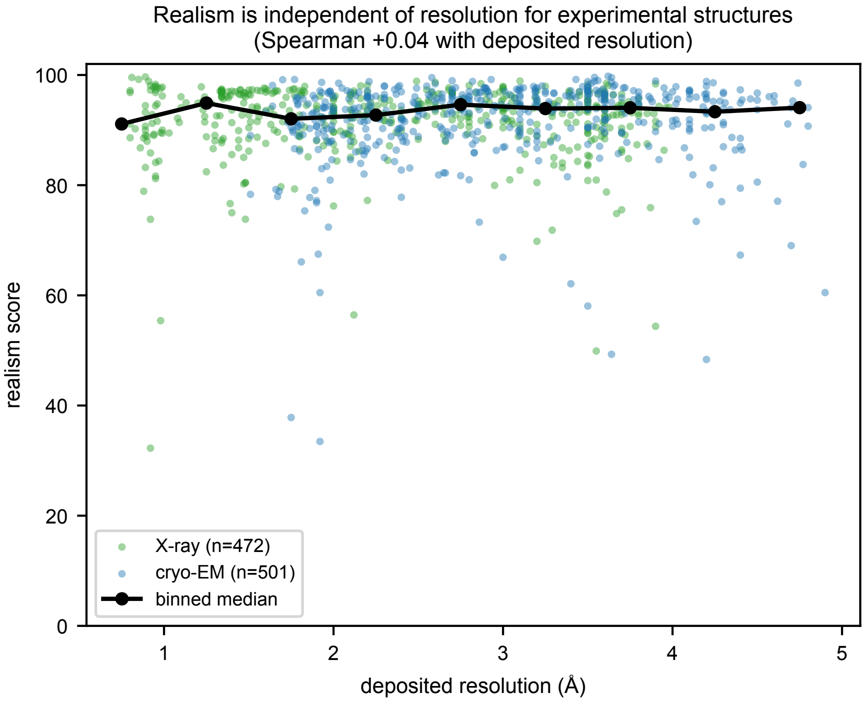


**Supplementary Figure 1. Realism is independent of resolution for experimental structures.** Realism score against deposited resolution for experimental X-ray and cryo-EM structures, with the binned median overlaid. Realism remains high and approximately constant across the full resolution range (Spearman 0.04 with deposited resolution), confirming that the score reflects experimental-like geometry rather than the amount of experimental detail.

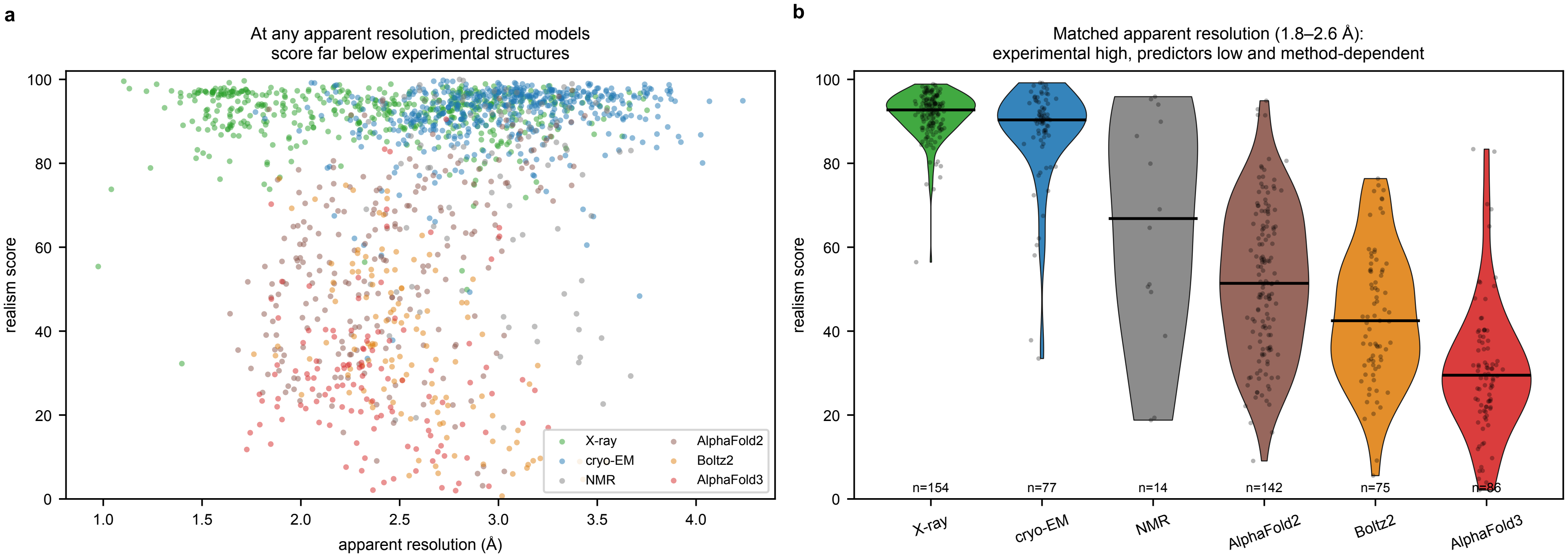


**Supplementary Figure 2. Predicted models score far below experimental structures at matched apparent resolution.** (a) Realism against apparent resolution across the benchmark, coloured by method: experimental structures (X-ray, cryo-EM, NMR) remain high across the apparent-resolution range, whereas AlphaFold2, AlphaFold3 and Boltz2 sit well below them even at favourable apparent resolutions. (b) Realism distributions within a matched apparent-resolution band (1.8 to 2.6 Å): experimental structures score around 92 while the predictors score substantially lower and method-dependently (AlphaFold2 51, Boltz2 42, AlphaFold3 29), showing the separation is not a resolution artefact.

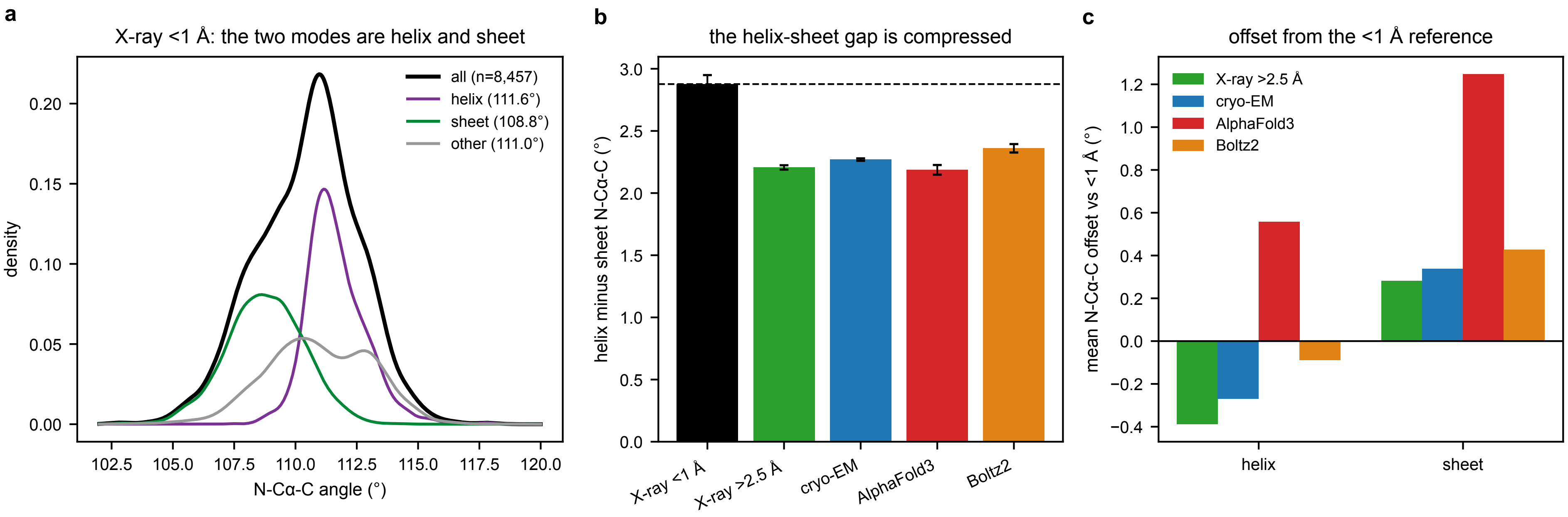


**Supplementary Figure 3. The N-Cα-C backbone angle by backbone conformation.** The two modes of the N-Cα-C angle distribution in Figure 2c are α-helical and β residues. Conformation is assigned from φ and ψ (α: −100° < φ < −30° and −80° < ψ < −10°; β: φ < −90° and ψ > 90°), and Gly and Pro are excluded as in Figure 2c. (a) In ultra-high-resolution (<1 Å) X-ray structures the mean angle is 111.6° for α-helical and 108.8° for β residues, a difference of 2.9°, reflecting the dependence of the angle on φ and ψ. (b) This helix-sheet difference is compressed to 2.2 to 2.4° in the lower-resolution X-ray, cryo-EM and predicted cohorts (error bars, 95% confidence interval). (c) Offset of the mean angle from the ultra-high-resolution reference: AlphaFold3 is shifted to larger angles in both conformations and most in β residues (+0.6° helix, +1.2° sheet), whereas Boltz2 stays close to the reference in helices (−0.1°).

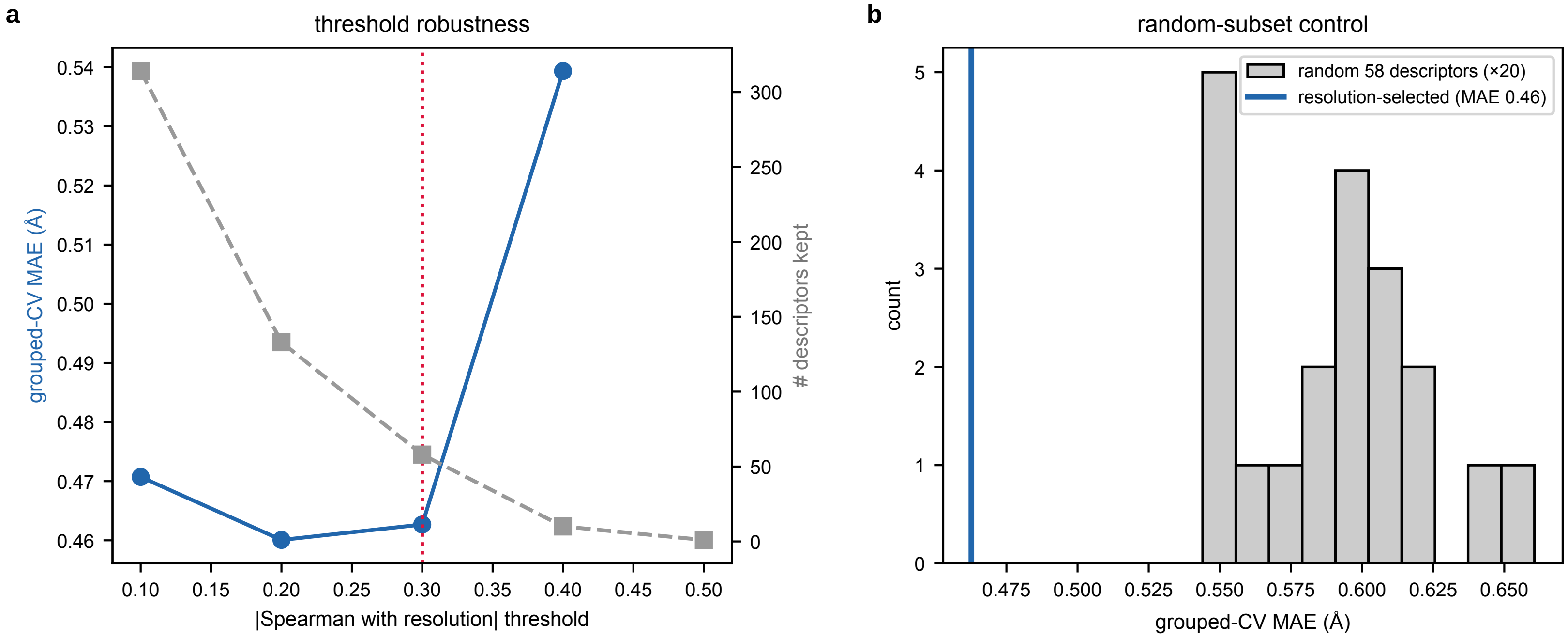


**Supplementary Figure 4. Robustness of the apparent-resolution descriptor selection.** (a) Grouped cross-validated mean absolute error as the correlation threshold for retaining resolution-scaling descriptors is swept from 0.1 to 0.5; with the selection repeated within each training fold, the error is stable through 0.3 (0.46 to 0.47 Å) and rises from 0.4 (0.54 Å); at 0.5 no descriptor passes in every training fold. (b) The resolution-selected descriptors (selected within each training fold) versus twenty random 58-descriptor draws from the size-invariant geometry panel: the selected set outperforms every random draw (random error 0.54 to 0.66 Å), so the signal is spread across many resolution-scaling features.

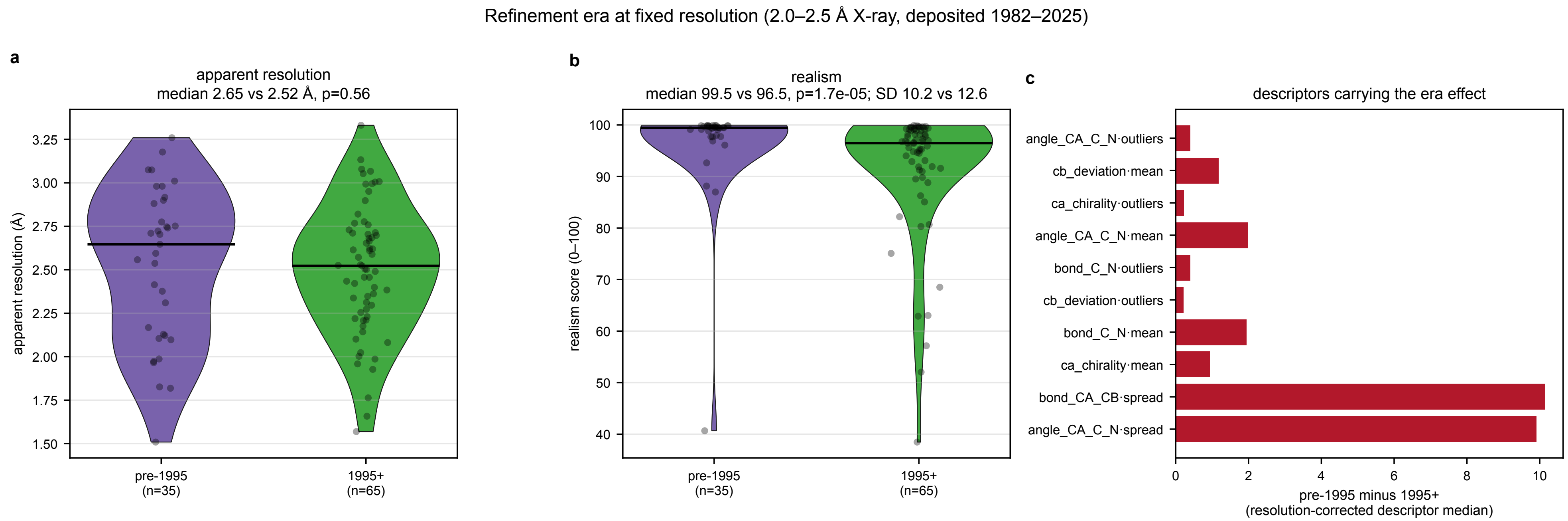


**Supplementary Figure 5. Apparent resolution and realism before and after 1995, at fixed resolution.** Crystallographic refinement has been progressively automated and re-standardised since the late 1980s, with the adoption of maximum-likelihood targets[42], improved stereochemical restraint libraries[21] and, later, conformation-dependent geometry restraints[18], which regularise model geometry at a given resolution, sometimes at the expense of experimentally supported variability[17]. We scored 100 X-ray structures at a fixed deposited resolution (2.0 to 2.5 Å): 35 deposited before 1995 and 65 from 1995 onward (deposition years 1982 to 2025), all re-featurised in a single pipeline run. (a) Apparent resolution is independent of era (median 2.65 versus 2.52 Å; Mann-Whitney $p = 0.56$; equal spread, Levene $p = 0.14$). (b) Realism is modestly lower for structures deposited from 1995 onward (median 96.5 for 1995-onward versus 99.5 for pre-1995, $p < 0.001$; Spearman with deposition year −0.40), with no significant difference in spread (standard deviation 12.6 versus 10.2, Levene $p = 0.09$). (c) The era-effect is confined to the restraint-sensitive local-geometry descriptors (backbone bond angles, peptide bond lengths, Cβ deviation, Cα chirality): structures deposited before 1995 carry more geometric scatter in exactly the quantities modern refinement homogenises. Apparent resolution is therefore unaffected by refinement era, while realism detects the known homogenisation of restrained local geometry; because this shifts modern structures slightly toward the idealised end, the effect is roughly an order of magnitude smaller than the experimental-versus-predicted separation on the realism axis (predicted medians 27 to 57 versus experimental 93 to 94) and leaves our conclusions unchanged, if anything making that contrast conservative.

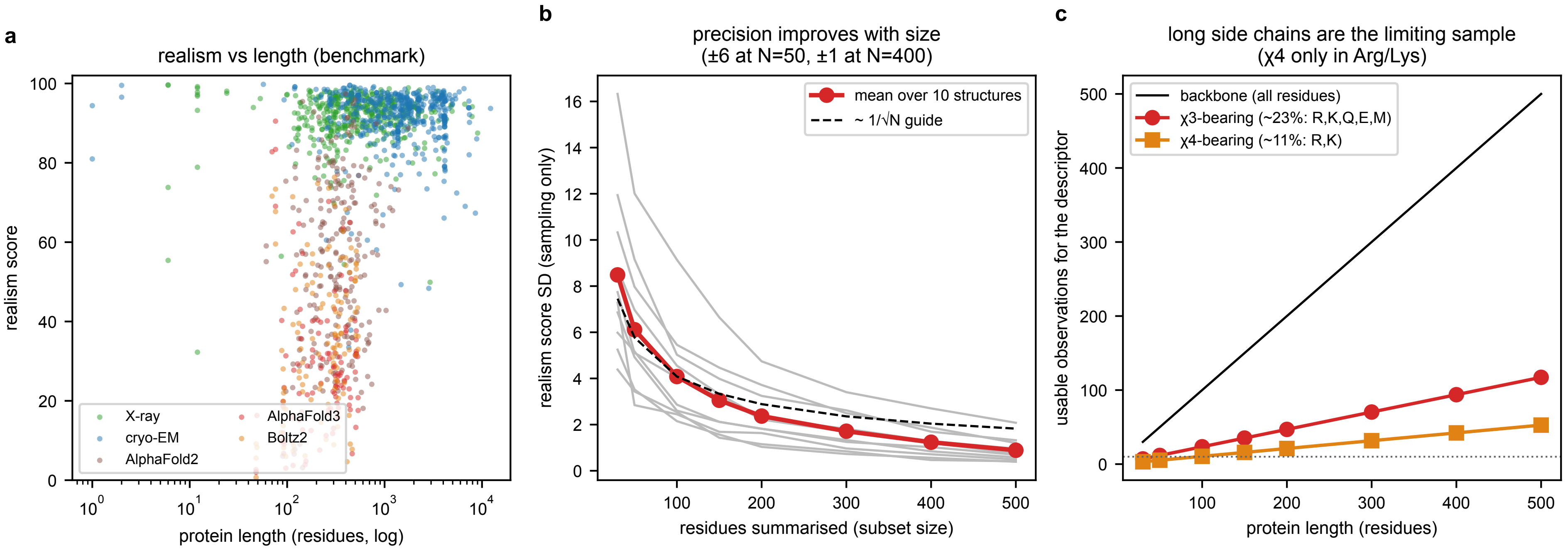


**Supplementary Figure 6. Protein size and score precision.** Both scores are summarised per residue across the whole structure, making them size-independent in expectation but not in precision. (a) Realism against protein length across the benchmark, coloured by method: the median shows no strong length trend, but the spread widens for small proteins. (b) Sampling uncertainty measured within single large structures: for ten X-ray structures of 614 to 663 residues, realism was recomputed from random residue subsets of increasing size (100 resamples each); the score standard deviation falls approximately as the inverse square root of the subset size, from about 6 points at 50 residues to about 1 at 400 or more, with negligible bias. (c) The limiting sample size is set by the long side chains: χ3 is defined for about 23% of residues (Arg, Lys, Gln, Glu, Met) and χ4 for only about 11% (Arg, Lys), so a 100-residue protein provides roughly ten χ4 observations.

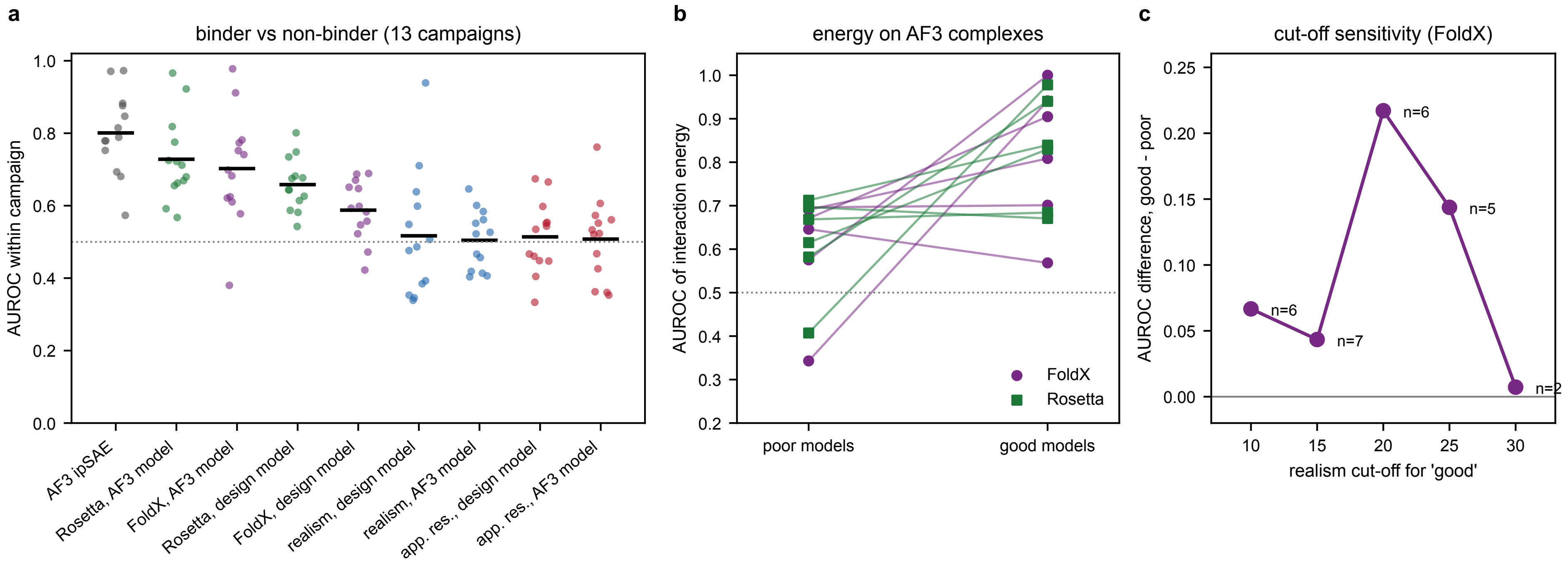


**Supplementary Figure 7. Model quality and the experimental success of de novo binders.** Data from a meta-analysis of 3,669 experimentally tested designs (394 binders)[24]. (a) Within-campaign AUROC for separating binders from non-binders in the 13 campaigns with at least five binders and five non-binders (points, one per campaign; bars, means), for the AlphaFold3 interface score ipSAE, the Rosetta and FoldX interaction energies computed on the AlphaFold3 and design models, and WhatRes realism and apparent resolution of the same complexes. (b) AUROC of the interaction energy on AlphaFold3 complexes rated poor or good by WhatRes (good: realism ≥ 20 and apparent resolution ≤ 3.0 Å), for the 6 campaigns with at least three binders and three non-binders in both groups; lines join the same campaign. (c) Mean AUROC difference between good and poor models for the FoldX energy, against the realism threshold; n, number of campaigns that can be tested.
